# Highly efficient synthetic CRISPR RNA/Cas9-based mutagenesis for rapid cardiovascular phenotypic screening in F0 zebrafish

**DOI:** 10.1101/2021.07.01.450753

**Authors:** Rachael E. Quick, Luke D. Buck, Sweta Parab, Zane R. Tolbert, Ryota L. Matsuoka

## Abstract

The zebrafish is a valuable vertebrate model to study cardiovascular formation and function due to the facile visualization and rapid development of the circulatory system in its externally growing embryos. Despite having distinct advantages, zebrafish have paralogs of many important genes, making reverse genetics approaches inefficient since generating animals bearing multiple gene mutations requires substantial efforts. Here, we present a simple and robust synthetic CRISPR RNA/Cas9-based mutagenesis approach for generating biallelic F0 zebrafish knockouts. Using a dual-guide synthetic CRISPR RNA/Cas9 ribonucleoprotein (dgRNP) system, we compared the efficiency of biallelic gene disruptions following the injections of one, two, and three dgRNPs per gene into the cytoplasm or yolk. We show that simultaneous cytoplasmic injections of three distinct dgRNPs per gene into one-cell stage embryos resulted in the most efficient and consistent biallelic gene disruptions. Importantly, this triple dgRNP approach enables efficient inactivation of cell autonomous and cell non-autonomous gene function, likely due to the low mosaicism of biallelic disruptions. In support of this finding, we provide evidence that the F0 animals generated by this method fully phenocopied the endothelial and peri-vascular defects observed in corresponding stable mutant homozygotes. Moreover, this approach faithfully recapitulated the trunk vessel phenotypes resulting from the genetic interaction between two *vegfr2* zebrafish paralogs. Mechanistically, investigation of genome editing and mRNA decay indicates that the combined mutagenic actions of three dgRNPs per gene lead to an increased probability of frameshift mutations, enabling efficient biallelic gene disruptions. Therefore, our approach offers a highly robust genetic platform to quickly assess novel and redundant gene function in F0 zebrafish.

## INTRODUCTION

The zebrafish model has become an increasingly popular vertebrate model to study dynamic tissue morphogenesis and cell physiology in living multicellular organisms. In particular, this model organism has provided a powerful screening and live imaging platform to study the mechanisms of organ growth, regeneration, and physiology due to their high fecundity, optical clarity, and regenerative capacity. As one of the first organ systems that develop during embryogenesis, the cardiovascular system has received much attention and its biological studies greatly benefited from research that utilized this model system [1–4]. Findings and insights from zebrafish studies have substantially contributed to expand our understanding of the vertebrate cardiovascular system in health and disease.

The advent of genome editing tools such as zinc-finger nucleases (ZFNs), transcription activator-like effector nucleases (TALENs), and CRISPR/Cas9 accelerates zebrafish genetic studies using reverse genetics approaches by facilitating the generation of targeted zebrafish mutants [5–11]. However, accumulating evidence has suggested that, due to the presence of duplicated genes or adapting genes whose responses are triggered by transcriptional adaptation, knocking out a single gene is not always sufficient to identify gene function through a phenotypic analysis of zebrafish mutants [12–15]. Since generating zebrafish that carry multiple mutated genes requires time-consuming, labor-intensive genetic crosses and subsequent genotyping, there is an increasing demand and necessity for a reliable, efficient method to quickly test the gene of interest and potential genetic redundancy in F0 animals.

Many studies have reported efficient methods for generating F0 zebrafish mutants using *in vitro*-transcribed guide RNA (gRNA) [16–22], or more recently, using a dual-guide CRISPR/Cas9 ribonucleoprotein (dgRNP) system that utilizes a chemically synthesized duplex of CRISPR RNA (crRNA) and trans-activating crRNA (tracrRNA) [23–30]. However, there is currently no integrated, standardized approach that is well established in the community, making it difficult to select which method to utilize based on pre-existing and emerging information in the rapidly evolving field. Current evidence suggests that synthetic crRNAs perfectly matched to target sequences achieve much more efficient target cleavage than that which *in vitro*-transcribed gRNAs carrying mismatched nucleotides are able to achieve in zebrafish [23]. This finding reported by Hoshijima *et al*. in late 2019 has allowed the broader use of a synthetic crRNA-based dgRNP method as a potentially more efficient genome editing approach. However, despite the higher efficacy of synthetic crRNAs at cleaving target sequences in general, their ability to induce monoallelic or biallelic gene inactivation in F0 animals depends on the target sequences of individual crRNAs, since variable efficiency of crRNAs in gene editing has been reported [29]. To circumvent this variable effect of individual crRNAs, a recent study injected an increasing number of dgRNPs per gene and showed enhanced efficiency of biallelic inactivation [24]. This study by Kroll *et al*. injected 1-4 dgRNPs per gene by targeting two pigmentation genes and showed that injections of three different dgRNPs per gene achieved more consistent biallelic disruptions, converting over 90% of injected embryos into F0 knockouts [24]. However, this study utilized exclusively yolk injections and eye pigmentation scoring as the phenotypic readout for this comparison, leaving open the question of injection site choice and optimal number of dgRNPs per gene in broader biological contexts. Further investigations into these questions will help establish a simple method that ensures maximal efficiency of generating biallelic F0 knockouts without requiring strict crRNA selections.

Here we addressed this issue by integrating some key observations from these prior studies. We specifically focused on assessing the ability of one, two, and three dgRNPs per gene to induce biallelic knockout phenotypes via cytoplasmic or yolk injections. We targeted multiple genetic loci critical for vascular development – phenotypic readouts that have not been extensively tested using the dgRNP system. Our in-depth quantitative comparisons using multiple phenotypic readouts demonstrate that simultaneous cytoplasmic injections of three different dgRNPs per gene outperformed all other tested injection conditions and resulted in the most efficient and consistent biallelic gene disruptions.

## RESULTS

### Cytoplasmic injections of three dgRNPs per gene increase the efficiency of generating biallelic F0 knockouts in zebrafish

Like in mammals, blood and lymphatic vessel development in zebrafish relies heavily on the coordinated actions of vascular endothelial growth factor (Vegf) signalling through its cognate Vegf receptors (Vegfrs). In the zebrafish genome, two *vegfr2* paralogs exist – *kdrl* and *kdr* [31], making a Vegf receptor family consisting of Flt1, Kdrl, Kdr, and Flt4. Several mutations in *kdrl* have been characterized in regards to their developmental vascular phenotypes, with *kdrl^um19^* mutants exhibiting a stalling in the sprouting of arterial intersegmental vessels (aISVs) from the dorsal aorta. *kdrl^um19^* mutants harbor a 4 base pair deletion in exon 2, resulting in a truncated extracellular domain without the receptor tyrosine kinase and transmembrane domains, which is expected to be a null mutant [32].

By analyzing progeny derived from *kdrl*^um19^ heterozygous adult incrosses (Figure 1A), we observed that *kdrl*^um19^ homozygous embryos displayed stalled aISV growth (Figures 1D, 1H), with some reaching the dorsal side of the trunk at 32 hours post fertilization (hpf) as previously reported [32]. In contrast, *kdrl*^um19^ heterozygotes exhibited ISV lengths equivalent to those seen in their WT siblings (Figures 1B, 1C, 1H). This stalled aISV phenotype observed in *kdrl*^um19^ homozygotes persisted at later developmental stages, up to at least 55 hpf (Figures 1E–G). Thus, we chose to utilize this robust and highly penetrant phenotype as a readout to test our approach in assessing monoallelic or biallelic disruption of this gene.

**Figure 1.**
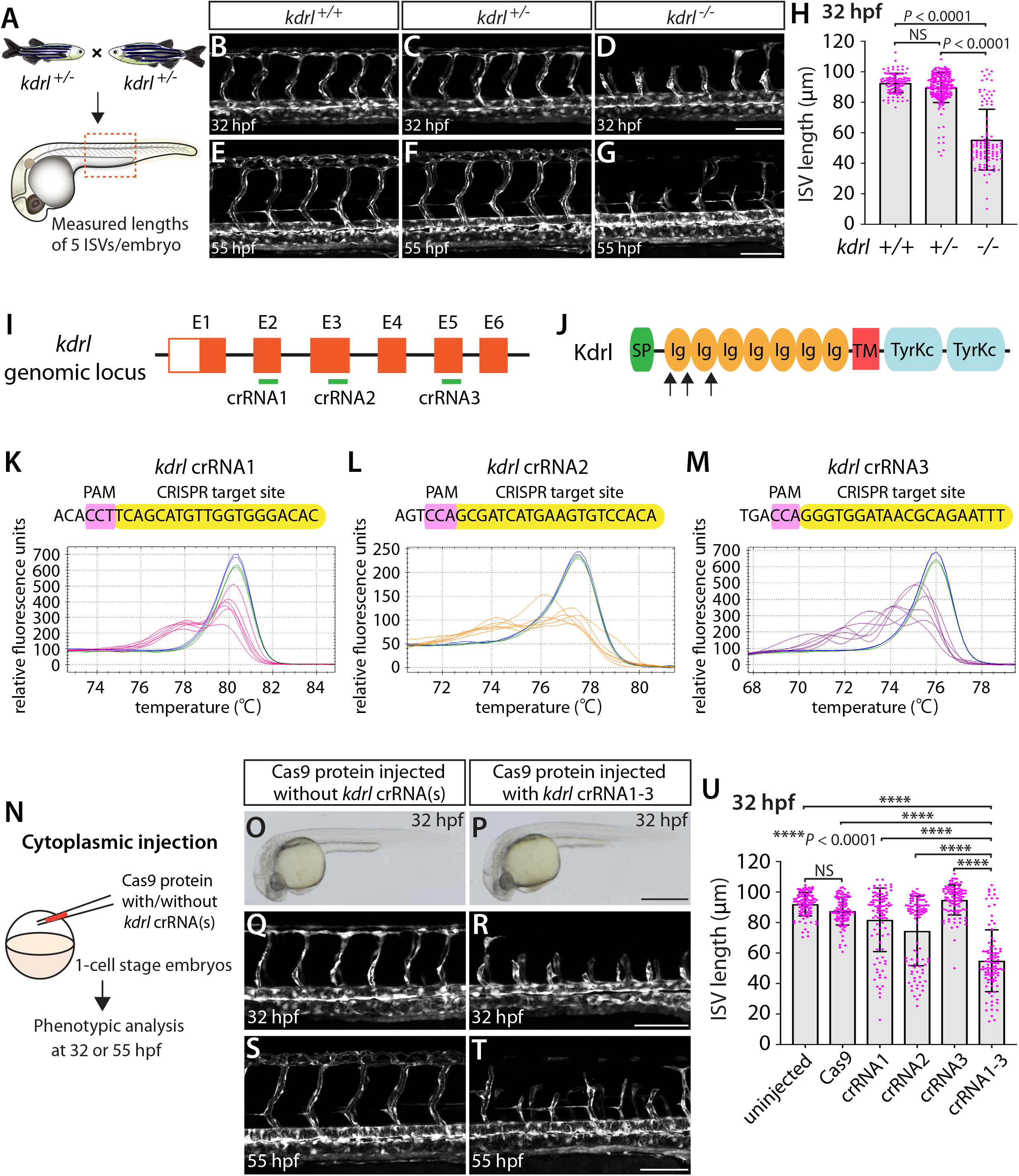
Embryos injected with the triple dgRNPs targeting *kdrl* exhibited aISV defects similar to those observed in *kdrl^−/−^* fish. (**A**) Experimental setup for the phenotypic analysis presented in panels (**B**–**H**). Adult *kdrl^+/−^* fish were incrossed, and progeny generated from the crosses were imaged to quantify the lengths of their arterial intersegmental vessels (aISVs) at 32 hpf. (**B**–**G**) Lateral views of *kdrl^+/+^* (**B**, **E**), *kdrl^+/−^* (**C**, **F**), and *kdrl^−/−^* (**D**, **G**) trunk vasculature visualized by *Tg(kdrl:*EGFP) expression at 32 (**B**–**D**) and 55 (**E**–**G**) hpf. (**H**) Quantification of aISV lengths in *kdrl^+/+^*, *kdrl^+/−^*, and *kdrl^−/−^* embryos at 32 hpf (n=23 for *kdrl^+/+^*, n=45 for *kdrl^+/−^*, and n=21 for *kdrl^−/−^* fish). Lengths of 5 aISVs per embryo were measured. NS: not significant. (**I**) Three synthetic CRISPR RNAs (crRNAs) were designed to target sequences within exon 2 (E2), E3, and E5 on the *kdrl* genomic locus. (**J**) Predicted domain structure of zebrafish Kdrl. Kdrl consists of a signal peptide (SP), seven immunoglobulin-like domains (Ig), a transmembrane domain (TM), and two tyrosine kinase domains (TyrKc). Arrows indicate the approximate positions of the protein sequences corresponding to the target sequences of the three designed crRNAs. (**K**–**M**) High-resolution melt analysis (HRMA) used to validate the efficacy of the three designed crRNAs. The melting curves of 6 independent embryos injected with the dgRNP complex containing *kdrl* crRNA1 (**K**, **pink**), crRNA2 (**L**, **orange**), or crRNA3 (**M**, **purple**) are presented. The melting curves of 2 independent, uninjected (**K**–**M**, **blue**) and Cas9 protein-injected (**K**–**M**, **green**) sibling embryos are also presented for comparison for each crRNA. Injection of each dgRNP cocktail disrupted the corresponding target genomic sequences. (**N**) Experimental workflow of the microinjection experiments for panels (**O**–**U**). Injection cocktails containing Cas9 protein with and without the individual or the three *kdrl* crRNA:tracrRNA duplexes were injected into the cytoplasm of one-cell stage *Tg(kdrl:EGFP)* embryos. Injected *Tg(kdrl:EGFP)* progeny were analyzed for aISV formation at 32 and 55 hpf. (**O** and **P**) Brightfield images of the 32 hpf embryos injected with Cas9 protein with (**P**) or without (**O**) the three *kdrl* crRNA:tracrRNA duplexes (crRNA1-3). (**Q**–**T**) Lateral trunk views of the 32 (**Q**, **R**) and 55 (**S**, **T**) hpf *Tg(kdrl:EGFP)* fish injected with Cas9 protein with (**R**, **T**) or without (**Q**, **S**) the three *kdrl* crRNA1-3. While aISVs in Cas9-injected embryos fused at the dorsal side of the trunk to form the dorsal longitudinal anastomotic vessel by 32 hpf (**Q**), a vast majority of aISVs in the embryos injected with the triple *kdrl* dgRNPs failed to reach the dorsal side of the trunk (**R**). This aISV stalling phenotype was observed at later developmental stages, including at 55 hpf (**T**). (**U**) Quantification of aISV lengths of 32 hpf embryos of indicated treatment (n=20 fish for each treatment group; 5 aISV lengths per embryo measured). Average aISV lengths of the triple dgRNPs-injected embryos closely resembled those of *kdrl^−/−^* fish. A one-way analysis of variance (ANOVA) followed by Tukey’s post-hoc test was used to calculate P values for panels (**H**, **U**). Scale bars: 50 µm in **D**, **G**, **R**, and **T**; 500 µm in **P**.

We first tested the efficacy of each designed *kdrl* crRNA. To this aim, we injected them individually into the cytoplasm of one-cell stage embryos and then carried out high-resolution melt analysis (HRMA) at 24 hpf (Figures 1I–M). We designed three different crRNAs (Figure 1I), all of which targeted the exons of *kdrl* that encode part of the 1st and 2nd immunoglobulin-like domains in the extracellular domain (Figure 1J). We expected these designed crRNAs to induce frameshift mutations that truncate Kdrl in the extracellular domain, resulting in a truncated protein with no receptor function. We validated that all crRNAs were capable of disrupting the corresponding target genomic sequences as revealed by HRMA (Figures 1K–M). We injected the three crRNAs individually, in the form of dgRNP complexes, into an endothelial-cell specific *Tg(kdrl:EGFP)* reporter line to compare their effect on aISV growth at 32 hpf (Figure 1N). This is the same developmental stage at which we analyzed embryos from *kdrl*^um19^ heterozygous incrosses. For phenotypic quantifications throughout this study, we chose to analyze only the embryos/larvae that did not display obvious gross morphological defects regardless of being injected or uninjected. Also, we injected Cas9 protein without a duplex of crRNAs and tracrRNAs as a control. We noted that the aISV lengths in these injected embryos varied considerably between the injection groups and also within each injection group. For example, the average aISV lengths (94.8 µm) of fish injected with the *kdrl* crRNA3-containing dgRNP complex closely matched those of the uninjected (92 µm) and Cas9-injected (87.5 µm) control siblings (Figure 1U). This result indicates that this particular dgRNP complex was not efficient at generating biallelic *kdrl* disruptions. In contrast, injections of the dgRNP complex containing crRNA1 or crRNA2 led to a broader range of aISV lengths (Figure 1U), including the stalled aISV phenotype observed in *kdrl^−/−^* embryos. However, despite this phenotype, overall average aISV lengths (81.8 and 74.5 µm for crRNA1 and crRNA2, respectively) were not comparable to those in *kdrl^−/−^* embryos (55.5 µm), suggesting that injections of individual dgRNP complexes were not efficient at generating biallelic *kdrl* knockouts.

Previous studies showed that increasing the number of injected *in vitro*-transcribed guide RNAs per gene enhances the efficiency of gene disruption [16]. Based on these data, we next asked whether a simultaneous cytoplasmic injection of the three dgRNP complexes increases the efficiency of *kdrl* gene disruptions. At the one-cell stage, we injected a mixture containing all three dgRNP complexes, each present at the same concentration of crRNAs used for their individual injections. We maintained the equal concentration of Cas9 protein across the individual, combined, and control injection cocktails for quantitative comparisons. We found that a simultaneous cytoplasmic injection of the three dgRNP complexes dramatically increased the number of embryos exhibiting the stalled aISV phenotype, as examined at 32 and 55 hpf (Figures 1Q–U) without showing apparent gross morphological defects (Figures 1O, 1P). When quantified for aISV lengths, we noted that the average aISV lengths (54.9 µm) in the triple dgRNPs-injected embryos closely resembled those observed in *kdrl^−/−^* embryos (Figure 1U). These results demonstrate that cytoplasmic injection of three dgRNPs per gene dramatically enhanced the efficiency of generating biallelic F0 *kdrl* knockouts that recapitulate the early embryonic trunk vascular defect seen in stable mutant homozygotes.

### Cytoplasmic injections of three unique *flt4* or *ccbe1* dgRNPs led to the complete loss of a meningeal perivascular cell type as observed in corresponding stable homozygous mutants

Next, we compared single and triple dgRNP injection approaches by examining a developmental process that occurs at later embryonic/larval stages and that involves another vascular cell type. Fluorescent granular perithelial cells (FGPs) – also called Mato cells [33], meningeal mural lymphatic endothelial cells [34], or brain lymphatic endothelial cells [35] – are a perivascular cell type that resides within the brain meninges. This cell type has been suggested to regulate meningeal angiogenesis and guide vascular regeneration after cerebrovascular injury in zebrafish [34, 36]. We hereafter refer to this cell type as FGPs. Zebrafish FGPs were previously shown to emerge from the optic choroidal vascular plexus at 2.5 dpf and migrate along the bilateral mesencephalic veins over the optic tectum by 4 dpf [33–35]. FGPs in zebrafish exhibit molecular signatures similar to those of lymphatic endothelial cells (LECs) and, like LECs, the formation of this cell type depends on the Vegfc/Vegfd/Flt4 signaling axis [34, 35]. Adult *flt4^hu4602^* mutants display a greatly reduced number of FGPs, and larvae injected with a morpholino targeting *flt4* exhibit completely absent FGPs at 5 dpf [34].

Since the *flt4^hu4602^* point mutation in the kinase domain is thought to be hypomorphic, we employed fish harboring a 2 base pair deletion in exon 4, the *flt4^um131^* mutation, which truncates Flt4 in the extracellular domain, thus resulting in a null mutant [12, 37]. To examine FGP formation in *flt4^um131^* mutants, we incrossed *flt4^um131^* heterozygous adults and quantified the number of FGPs that form the bilateral loops over the optic tectum in *flt4^+/+^*, *flt4^+/−^*, and *flt4^−/−^* larvae at 6 dpf (Figures 2A–E) – 2 days after FGP migration over the optic tectum is complete in WT. To quantify the number of FGPs, we employed *Tg(fli1:nEGFP);Tg(lyve1:DsRed)* double transgenic lines, which label all endothelial cell nuclei in EGFP and mark lymphatic cell lineages, including FGPs, in DsRed. We counted the number of *Tg(fli1:nEGFP);Tg(lyve1:DsRed)* double-positive cells that comprised the bilateral loops over the optic tectum within the defined areas (Figure S1). We noted that, despite the broader range in the number of FGPs observed in *flt4^+/−^* larvae compared to their *flt4^+/+^* counterparts, these two genotypes did not display a significant difference in cell numbers at 6 dpf (Figures 2B, 2C, 2E). However, all *flt4^−/−^* larvae examined at 6 dpf completely lacked FGPs (Figures 2D, 2E). We used this clear FGP phenotype as another readout to assess the efficiency of single and triple dgRNP injections in generating biallelic disruption of the *flt4* gene.

**Figure 2.**
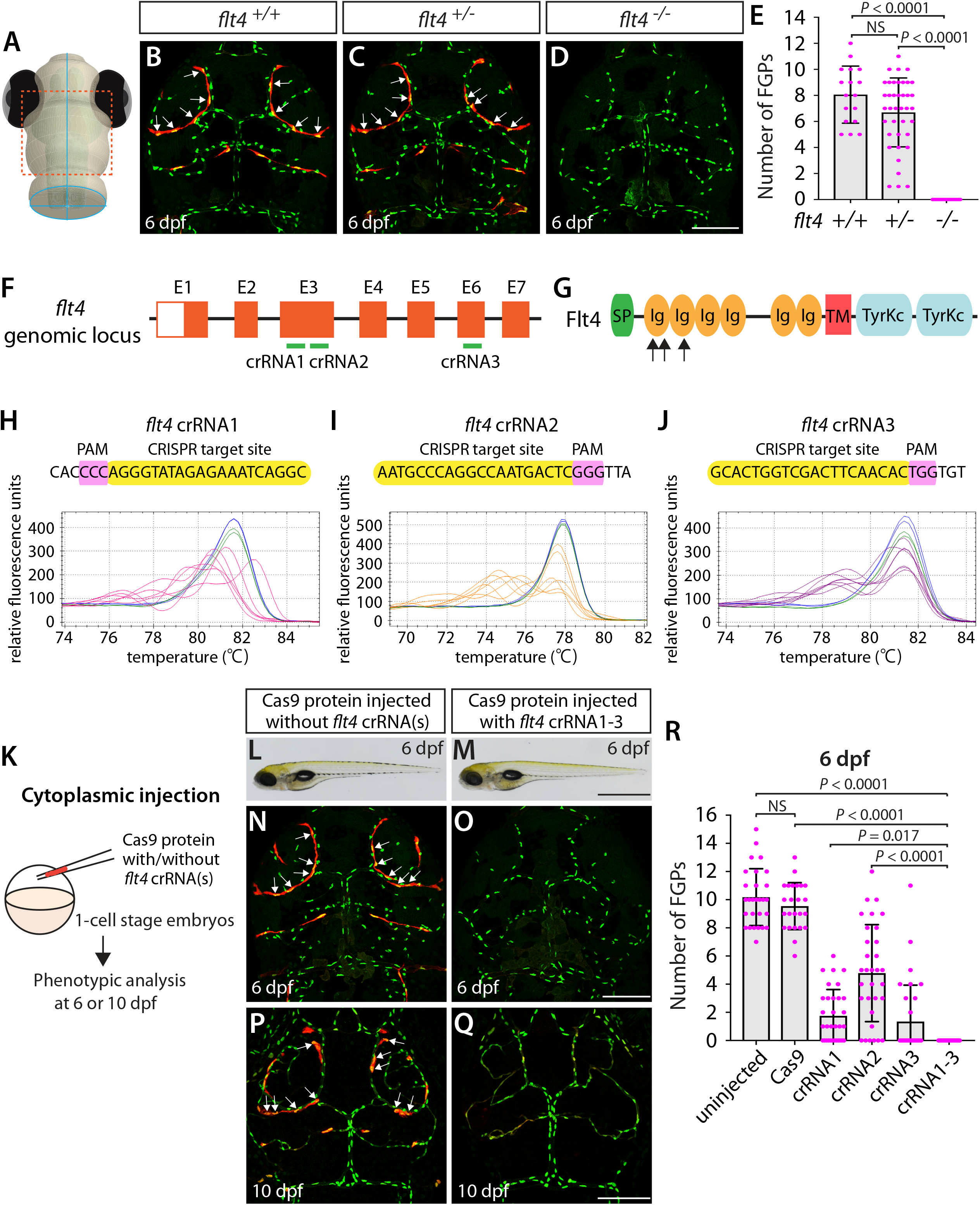
Larvae injected with three unique *flt4* dgRNPs phenocopied the complete loss of brain meningeal perivascular FGPs observed in *flt4^−/−^* fish. (**A**) Schematic representation of the dorsal view of the zebrafish larval head. The boxed area indicates the approximate region where the confocal images of panels (**B**–**D**) were captured. (**B**–**D**) Dorsal head views of 6 dpf *flt4^+/+^*, *flt4^+/−^*, and *flt4^−/−^* larvae carrying *Tg(fli1:nEGFP)* and *Tg(lyve1:DsRed)* transgenes. While *flt4^+/+^* and *flt4^+/−^* larvae formed *Tg(fli1:*nEGFP*);Tg(lyve1:*DsRed)-double positive FGPs in the dorsal meningeal surfaces over the optic tectum (arrows, **B** and **C**), *flt4^−/−^* larvae completely lacked FGPs (**D**). (**E**) Quantification of FGPs over the optic tectum for each genotype at 6 dpf (n=17 for *flt4^+/+^*, n=39 for *flt4^+/−^*, and n=21 for *flt4^−/−^* fish). (**F**) Three synthetic crRNAs were designed to target sequences within exon 3 (E3) and E6 on the *flt4* genomic locus. (**G**) Predicted domain structure of zebrafish Flt4. Flt4 consists of a signal peptide (SP), six immunoglobulin-like domains (Ig), a transmembrane domain (TM), and two tyrosine kinase domains (TyrKc). Arrows indicate the approximate positions of the protein sequences corresponding to the target sequences of the three designed crRNAs. (**H**–**J**) HRMA used to validate the efficacy of the three designed crRNAs. The melting curves of 6 independent embryos injected with the dgRNP complex containing *flt4* crRNA1 (**H**, **pink**), crRNA2 (**I**, **orange**), or crRNA3 (**J**, **purple**) are presented. The melting curves of 2 independent, uninjected (**H**–**J**, **blue**) and Cas9-injected (**H**–**J**, **green**) sibling embryos are also presented for comparison for each crRNA. Injection of each dgRNP cocktail disrupted the corresponding target genomic sequences. (**K**) Experimental workflow of the microinjection experiments for panels (**L**–**R**). Injection cocktails containing Cas9 protein with and without the individual or the three *flt4* crRNA:tracrRNA duplexes were injected into the cytoplasm of one-cell stage *Tg(fli1:nEGFP);Tg(lyve1:DsRed)* embryos. Injected progeny were analyzed at 6 and 10 dpf for FGP formation over the optic tectum. (**L** and **M**) Brightfield images of the 6 dpf larvae injected with Cas9 protein with (**M**) or without (**L**) the three *flt4* crRNA:tracrRNA duplexes (crRNA1-3). (**N**–**Q**) Dorsal head views of the 6 (**N**, **O**) and 10 (**P**, **Q**) dpf *Tg(fli1:nEGFP);Tg(lyve1:DsRed)* larvae injected with Cas9 protein with (**O**, **Q**) or without (**N**, **P**) the three *flt4* crRNA1-3. While larvae injected with Cas9 alone formed FGPs over the optic tectum at 6 and 10 dpf (arrows, **N** and **P**), those injected with the three *flt4* dgRNPs displayed a complete lack of the FGPs at both stages (**O** and **Q**). (**R**) Quantification of FGPs over the optic tectum at 6 dpf (n=28 for uninjected; n=24 for Cas9 controls; n=32 for crRNA1, crRNA2, and crRNA3; and n=35 for crRNA1-3). Larvae injected with all three *flt4* dgRNPs failed to form FGPs. Fish injected with the individual *flt4* dgRNPs displayed varying degrees of FGP formation. A one-way ANOVA followed by Tukey’s post-hoc test was used to calculate P values for panels (**E**, **R**). Scale bars: 50 µm in **D**, **O**, and **Q**; 1 mm in **M**.

Similarly to how we validated the *kdrl* crRNAs, we examined the efficacy of each *flt4* crRNA by injecting them individually into one-cell stage embryos and then performing HRMA at 24 hpf (Figures 2F–J). We designed three different crRNAs (Figure 2F), all of which were targeted to the exons of *flt4* that encode part of the 1st and 2nd immunoglobulin-like domains in the extracellular domain (Figure 2G). We thus expected these designed crRNAs to cause frameshift mutations that truncate Flt4 in the extracellular domain, resulting in a truncated protein with no receptor function. These three crRNAs were able to disrupt the corresponding target genomic sequences as revealed by HRMA (Figures 2H–J). We injected them in the form of dgRNPs individually, or as a mixture, into *Tg(fli1:nEGFP);Tg(lyve1:DsRed)* double transgenic lines to compare their effect on FGP numbers over the optic tectum at 6 dpf (Figure 2K). This is the same developmental stage at which we analyzed larvae from *flt4^um131^* heterozygous incrosses. Under these experimental conditions, single dgRNP injections exhibited varied efficacy. Injections of the crRNA2-containing dgRNP complex alone showed broader ranges in FGP numbers than those observed in the uninjected and Cas9-injected control siblings (Figure 2R). Injections of the dgRNP complex containing either crRNA1 or crRNA3 led to an overall reduction of FGP numbers compared to the control groups (Figure 2R). Approximately 38%, 19%, and 72% of larvae injected with the individual dgRNP cocktail containing crRNA1, crRNA2, or crRNA3, respectively, failed to form FGPs at 6 dpf. In contrast, combined injections of the three dgRNPs led to a complete absence of FGPs without displaying gross morphological deficits except slight pericardial edema (Figures 2L, 2M). This “no FGP” phenotype was seen in all injected larvae examined at 6 dpf (Figures 2N, 2O, 2R) as observed in *flt4^−/−^* larvae. This phenotype persisted at later developmental time points, up to at least 10 dpf (Figures 2P, 2Q). These results demonstrate that simultaneous cytoplasmic injections of the triple dgRNP cocktail allowed for highly efficient generation of biallelic F0 *flt4* knockouts, resulting in a complete loss of perivascular FGPs over the optic tectum.

Using the same readout, we next targeted a separate gene encoding the secreted protein Ccbe1. This experiment combined with the *flt4* dgRNP injection results allowed us to investigate the ability of a triple dgRNP injection approach in disrupting cell autonomous and cell non-autonomous gene function. As previously reported [35], *ccbe1* homozygous, but not heterozygous, mutants displayed a complete loss of FGPs over the optic tectum at 6 dpf (Figures 3A–E, S2). We then injected the three validated crRNAs targeting *ccbe1* (Figures 3F–J) in the form of dgRNPs individually, or simultaneously, into the cytoplasm of one-cell stage *Tg(fli1:nEGFP);Tg(lyve1:DsRed)* embryos (Figure 3K). The phenotypic results were very similar to what we observed in larvae injected with single or the triple *flt4* dgRNPs. Injections of individual *ccbe1* dgRNPs led to variable FGP phenotypes regardless of the crRNAs. Approximately 60%, 40%, and 90% of larvae injected with the single dgRNP cocktail containing crRNA1, crRNA2, or crRNA3, respectively, failed to form FGPs at 6 dpf. This result is in contrast to the consistent, fully penetrant “no FGP” phenotype in larvae injected with the three combined dgRNPs (Figures 3L–R). These findings suggest that the approach of injecting three dgRNPs per gene enables biallelic disruptions of cell autonomous and cell non-autonomous gene function.

**Figure 3.**
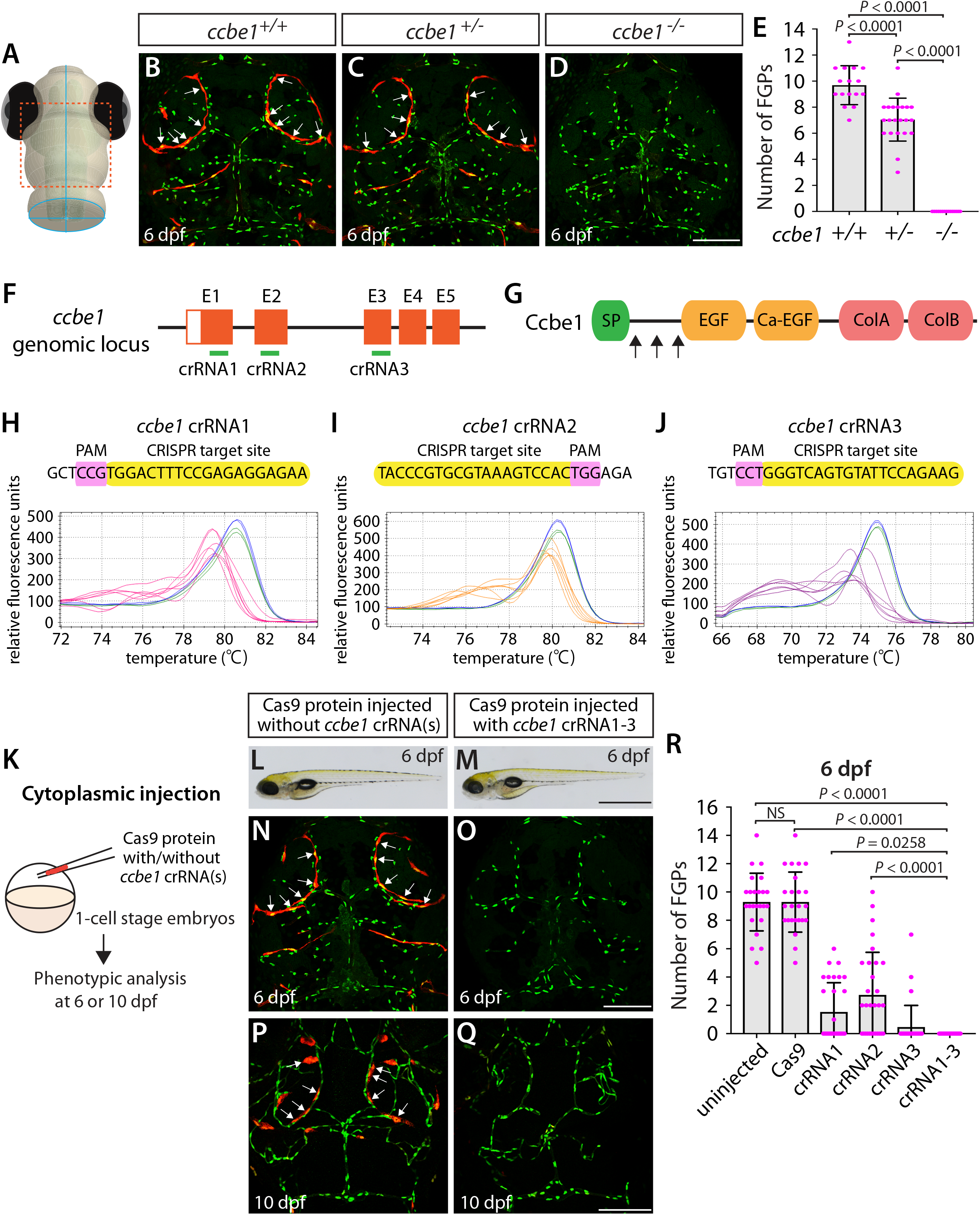
Larvae injected with three unique *ccbe1* dgRNPs phenocopied the complete loss of brain meningeal perivascular FGPs observed in *ccbe1^−/−^* fish. (**A**) Schematic representation of the dorsal view of the zebrafish larval head. The boxed area indicates the approximate region where the confocal images of panels (**B**–**D**) were captured. (**B**–**D**) Dorsal head views of 6 dpf *ccbe1^+/+^*, *ccbe1^+/−^*, and *ccbe1^−/−^* larvae carrying *Tg(fli1:nEGFP)* and *Tg(lyve1:DsRed)* transgenes. While *ccbe1^+/+^* and *ccbe1^+/−^* larvae formed *Tg(fli1:*nEGFP*);Tg(lyve1:*DsRed)-double positive FGPs in the dorsal meningeal surfaces over the optic tectum (arrows, **B** and **C**), *ccbe1^−/−^* larvae completely lacked FGPs (**D**). (**E**) Quantification of FGPs over the optic tectum for each genotype at 6 dpf (n=16 for *ccbe1^+/+^*, n=22 for *ccbe1^+/−^*, and n=15 for *ccbe1^−/−^* fish). (**F**) Three synthetic crRNAs were designed to target sequences within exon 1 (E1), E2 and E3 on the *ccbe1* genomic locus. (**G**) Predicted domain structure of zebrafish Ccbe1. Ccbe1 consists of a signal peptide (SP), an EGF domain (EGF), a calcium-binding EGF domain (Ca-EGF), and two collagen repeat domains (ColA and ColB). Arrows indicate the approximate positions of the protein sequences corresponding to the target sequences of the three designed crRNAs. (**H**–**J**) HRMA used to validate the efficacy of the three designed crRNAs. The melting curves of 6 independent embryos injected with the dgRNP complex containing *ccbe1* crRNA1 (**H**, **pink**), crRNA2 (**I**, **orange**), or crRNA3 (**J**, **purple**) are presented. The melting curves of 2 independent, uninjected (**H**– **J**, **blue**) and Cas9-injected (**H**–**J**, **green**) sibling embryos are also presented for comparison for each crRNA. Injection of each dgRNP cocktail disrupted the corresponding target genomic sequences. (**K**) Experimental workflow of the microinjection experiments for panels (**L**–**R**). Injection cocktails containing Cas9 protein with and without the individual or the three *ccbe1* crRNA:tracrRNA duplexes were injected into the cytoplasm of one-cell stage *Tg(fli1:nEGFP);Tg(lyve1:DsRed)* embryos. Injected progeny were analyzed at 6 and 10 dpf for FGP formation over the optic tectum. (**L** and **M**) Brightfield images of the 6 dpf larvae injected with Cas9 protein with (**M**) or without (**L**) the three *ccbe1* crRNA:tracrRNA duplexes (crRNA1-3). (**N**–**Q**) Dorsal head views of the 6 (**N**, **O**) and 10 (**P**, **Q**) dpf *Tg(fli1:nEGFP);Tg(lyve1:DsRed)* larvae injected with Cas9 protein with (**O**, **Q**) or without (**N**, **P**) the three *ccbe1* crRNA1-3. Larvae injected with Cas9 alone formed FGPs over the optic tectum at 6 and 10 dpf (arrows, **N** and **P**). However, fish injected with the three *ccbe1* dgRNPs completely lacked the FGPs at both stages (**O** and **Q**). (**R**) Quantification of FGPs over the optic tectum at 6 dpf (n=24 for uninjected and Cas9 controls; n=30 for crRNA1, crRNA2, and crRNA3; and n=35 for crRNA1-3). All larvae injected with the three *ccbe1* dgRNPs exhibited a complete loss of FGPs over the optic tectum. Fish injected with the individual *ccbe1* dgRNPs displayed varying degrees of FGP formation. A one-way ANOVA followed by Tukey’s post-hoc test was used to calculate P values for panels (**E**, **R**). Scale bars: 50 µm in **D**, **O**, and **Q**; 1 mm in **M**.

### Cytoplasmic injections of three unique *vegfab* dgRNPs led to the complete loss of the vertebral arteries as observed in *vegfab* stable homozygous mutants

A previous study showed that Vegfab is required for the formation of vertebral arteries (VTAs), which develop bilaterally in the ventrolateral locations adjacent to the spinal cord [38]. Endothelial cells (ECs) that form the VTAs sprout from the ISVs between 3 and 4 dpf and continue to extend through 7–8 dpf [38]. As reported in this prior study, we observed that VTAs were completely absent in all *vegfab^−/−^* larvae examined at 8 dpf (Figures 4A, 4D, 4E). The *vegfab^+/−^* larvae of the same age displayed an overall reduction in the number of ECs comprising the VTAs as compared to that observed in *vegfab^+/+^* fish (Figures 4B, 4C, 4E). Despite this lower EC number, nearly 94% of *vegfab^+/−^* larvae formed VTAs in contrast to the complete absence of VTAs in *vegfab^−/−^* fish. To quantify the exact number of ECs comprising VTAs, we employed *Tg(kdrl:EGFP);Tg(kdrl:NLS-mCherry)* double transgenic reporters in which blood ECs and their nuclei are marked in EGFP and mCherry, respectively. We used this clean phenotype as another readout to further test the approach of injecting single and triple dgRNPs targeted to the secreted angiogenic ligand Vegfab.

**Figure 4.**
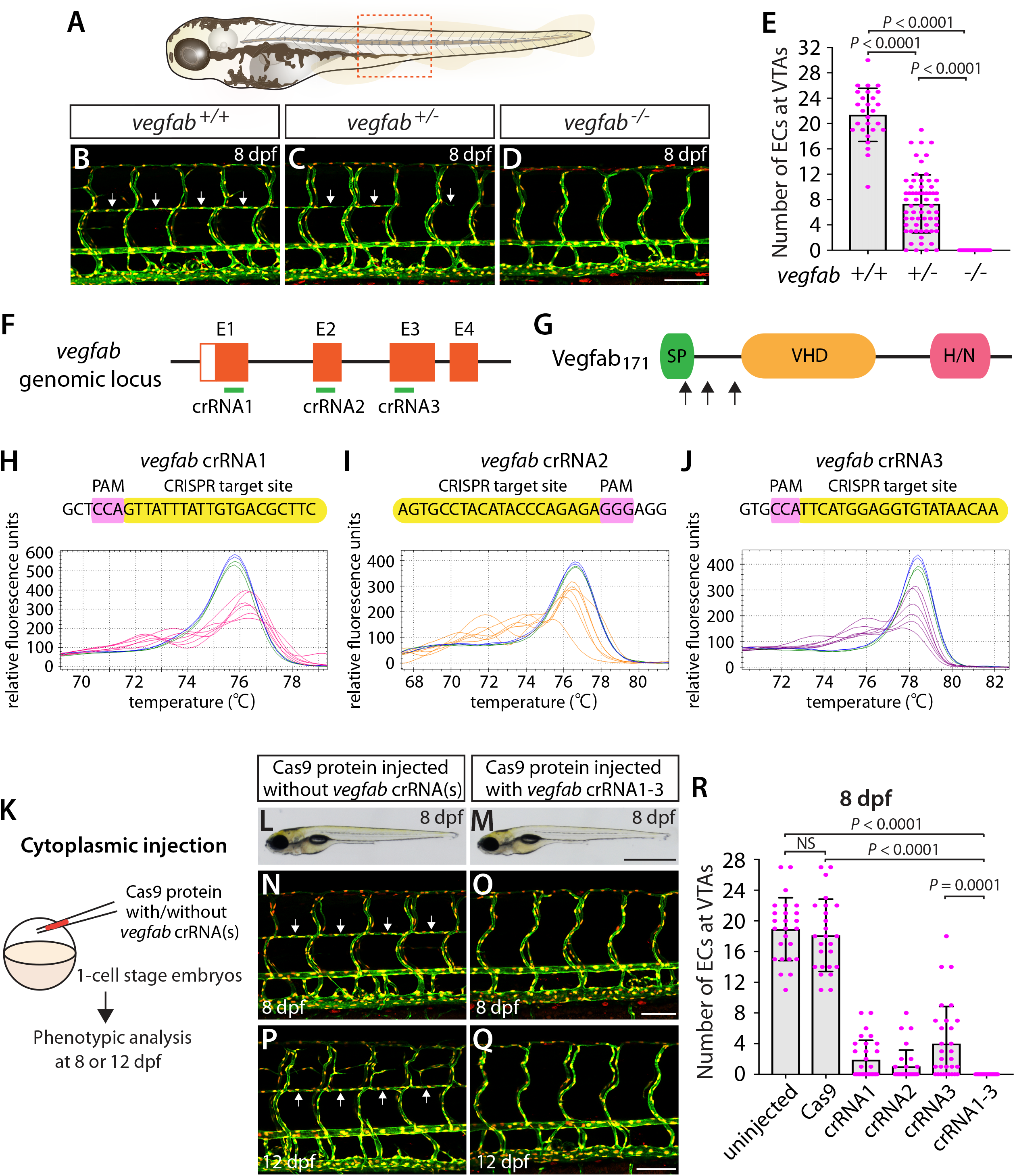
Larvae injected with three unique *vegfab* dgRNPs phenocopied the complete loss of VTAs observed in *vegfab^−/−^* fish. (**A**) Schematic representation of the lateral view of an 8 dpf zebrafish larva. The boxed area indicates the approximate trunk region where all VTA quantifications were performed. (**B**–**D**) Lateral trunk views of 8 dpf *vegfab^+/+^*, *vegfab^+/−^*, and *vegfab^−/−^* larvae carrying *Tg(kdrl:EGFP)* and *Tg(kdrl:NLS-mCherry)* transgenes. While *vegfab^+/+^* larvae formed the VTAs, which extended bilaterally along the ventrolateral sides of the spinal cord (**B**, arrows), *vegfab^−/−^* larvae formed no VTAs (**D**). Most *vegfab^+/−^* larvae exhibited partially forming VTAs at this stage (**C**, arrows). (**E**) Quantification of endothelial cells (ECs) at the VTAs within a 5 somite region per animal at 8 dpf (n=27 for *vegfab^+/+^*, n=64 for *vegfab^+/−^*, and n=35 for *vegfab^−/−^* fish). (**F**) Three synthetic crRNAs were designed to target sequences within exon 1 (E1), E2, and E3 on the *vegfab* genomic locus. (**G**) Predicted domain structure of zebrafish Vegfab_171_ isoform. Vegfab_171_ consists of a signal peptide (SP), a VEGF homology domain (VHD), and a heparin and neuropilin1 binding domain (H/N). Arrows indicate the approximate positions of the protein sequences corresponding to the target sequences of the three designed crRNAs. (**H**–**J**) HRMA used to validate the efficacy of the three designed crRNAs. The melting curves of 6 independent embryos injected with the dgRNP complex containing *vegfab* crRNA1 (**H**, **pink**), crRNA2 (**I**, **orange**), or crRNA3 (**J**, **purple**) are presented. The melting curves of 2 independent, uninjected (**H**– **J**, **blue**) and Cas9-injected (**H**–**J**, **green**) sibling embryos are also presented for comparison for each crRNA. Injection of each dgRNP cocktail disrupted the corresponding target genomic sequences. (**K**) Experimental workflow of the microinjection experiments for panels (**L**–**R**). Injection cocktails containing Cas9 protein with and without the individual or the three *vegfab* crRNA:tracrRNA duplexes were injected into the cytoplasm of one-cell stage *Tg(kdrl:EGFP);Tg(kdrl:NLS-mCherry)* embryos. Injected progeny were analyzed for VTA formation at 8 and 12 dpf. (**L** and **M**) Brightfield images of the 8 dpf larvae injected with Cas9 protein with (**M**) or without (**L**) the three *vegfab* crRNA:tracrRNA duplexes (crRNA1-3). (**N**–**Q**) Lateral trunk views of the 8 (**N**, **O**) and 12 (**P**, **Q**) dpf *Tg(kdrl:EGFP);Tg(kdrl:NLS-mCherry)* larvae injected with Cas9 protein with (**O**, **Q**) or without (**N**, **P**) the three *vegfab* crRNA1-3. While larvae injected with Cas9 alone formed most of the VTAs at 8 and 12 dpf (arrows, **N** and **P**), those injected with the three *vegfab* dgRNPs completely lacked VTAs at both stages (**O** and **Q**). (**R**) Quantification of ECs at the VTAs within a 5 somite region per animal at 8 dpf (n=25 for uninjected and Cas9 controls; n=30 for crRNA1, crRNA2, and crRNA3; and n=32 for crRNA1-3). All larvae injected with the three *vegfab* dgRNPs failed to form VTAs. Fish injected with the individual *vegfab* dgRNPs displayed varying degrees of VTA formation. A one-way ANOVA followed by Tukey’s post-hoc test was used to calculate P values for panels (**E**, **R**). Scale bars: 50 µm in **D**, **O**, and **Q**; 1 mm in **M**.

We designed three *vegfab* crRNAs by targeting exons that encode the N-terminal domain upstream of the VEGF homology domain critical for Vegf dimerization and function. These three crRNAs were all designed to target sequences upstream of the mutations harbored in the *vegfab^bns92^* allele (Figures 4F, 4G). We validated the efficacy of individual *vegfab* crRNAs by injecting each into one-cell stage embryos and then performing HRMA at 24 hpf (Figures 4H–J). We injected the three crRNAs in the form of dgRNPs individually, or simultaneously, into the cytoplasm of one-cell stage *Tg(kdrl:EGFP);Tg(kdrl:NLS-mCherry)* embryos to compare their effect on VTA formation at 8 dpf (Figure 4K). Uninjected and Cas9-injected fish displayed mostly formed VTAs at this stage (Figures 4L, 4N, 4R). We noted that each of the three dgRNPs targeting *vegfab* displayed varied efficiency at abrogating VTA formation. Approximately 53%, 70%, and 30% of larvae injected with the individual dgRNP cocktail containing crRNA1, crRNA2, or crRNA3, respectively, failed to form VTAs at 8 dpf. In contrast, all larvae injected with the triple dgRNPs exhibited a complete loss of VTAs at 8 dpf (Figures 4M, 4O, 4R). This “no VTA” phenotype persisted at later developmental stages, up to at least 12 dpf (Figures 4P, 4Q). These results provide another example of highly-efficient gene disruption via simultaneous cytoplasmic injection of three dgRNPs targeting the secreted protein, leading to a defect in trunk blood EC development.

### Yolk injections of three dgRNPs per gene allowed efficient generation of biallelic F0 knockouts but were less consistent than cytoplasmic injections

The dgRNP injection results we have presented thus far were all performed via cytoplasmic injections at the one-cell stage. This selection of injection sites is based on an original study that injected a RNP complex composed of synthetic crRNA, tracrRNA, and Cas9 protein into the cytoplasm for mutagenesis in zebrafish [23]. A recent study showed that yolk injections of dgRNP complexes are also highly effective [24]. However, these two studies did not conduct direct quantitative comparisons by injecting the same set of dgRNP complexes into different embryonic locations. Thus, we addressed this issue by performing yolk injections using the same phenotypic readouts and preparations of dgRNP complexes that were employed for the cytoplasmic injection experiments. This data allowed us to directly compare the two injection types.

First, we tested yolk injections of the three dgRNPs targeting *flt4* or *ccbe1* by examining FGP formation over the optic tectum at 6 dpf (Figure 5A). We noted that *flt4* and *ccbe1* triple dgRNP injections into the yolk were both highly efficient at abrogating FGP formation. All, except one, larvae (34 out of 35) injected with the three dgRNPs targeting *flt4* via the yolk displayed a complete lack of FGPs at 6 dpf (Figures 5B, 5C). On the other hand, we observed that 4 out of 35 larvae (approximately 11%) injected with the three dgRNPs targeting *ccbe1* via the yolk formed FGPs similar to Cas9-injected sibling controls, while the remaining larvae displayed no FGPs over the optic tectum. Since none of the larvae injected with the same three dgRNPs targeting *flt4* or *ccbe1* via the cytoplasm formed FGPs, cytoplasmic injections may lead to more consistent and efficient results than yolk injections.

**Figure 5.**
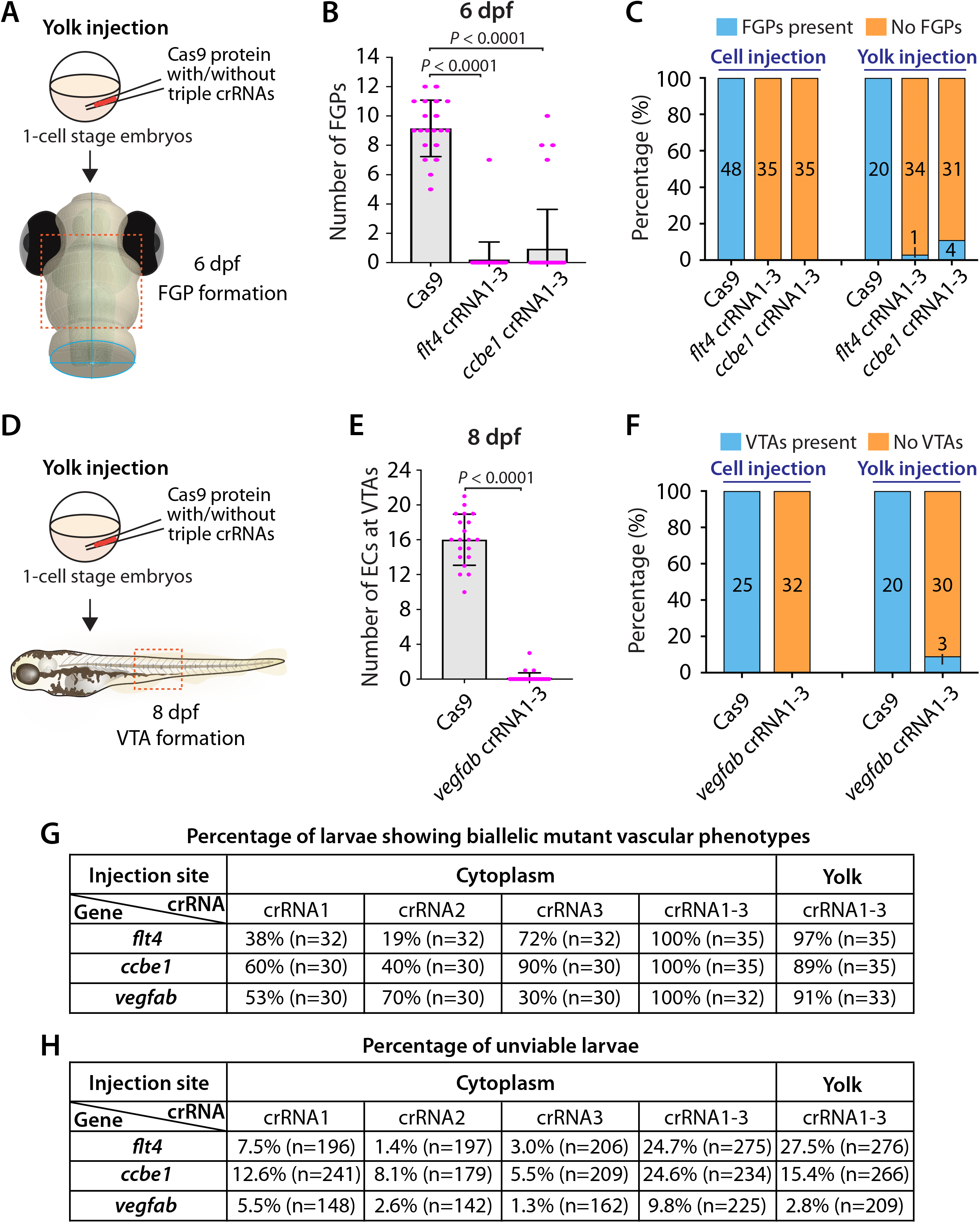
Yolk injections of three dgRNPs per gene allowed efficient generation of biallelic F0 knockouts but were less consistent than cytoplasmic injections. (**A**) Experimental workflow of the yolk microinjection experiments for panel (**B**). Injection cocktails containing Cas9 protein with and without the three *flt4* or *ccbe1* crRNA:tracrRNA duplexes were injected into the yolk of one-cell stage *Tg(fli1:nEGFP);Tg(lyve1:DsRed)* embryos. Injected progeny were analyzed at 6 dpf for FGP formation over the optic tectum. (**B**) Quantification of FGPs over the optic tectum at 6 dpf after yolk injection of Cas9 protein with and without the three *flt4* or *ccbe1* crRNA:tracrRNA duplexes (n=20 for Cas9 controls, n=35 for *flt4* crRNA1-3 and *ccbe1* crRNA1-3). All *flt4* dgRNPs-injected larvae except one failed to form FGPs over the optic tectum. Approximately 89% of *ccbe1* dgRNPs-injected larvae completely lacked FGPs, but the remaining fish formed FGPs in a manner comparable to Cas9-injected controls. A one-way ANOVA followed by Dunnett’s post-hoc test was used to calculate P values. (**C**) Percentage of 6 dpf larvae of indicated treatment with and without FGPs over the optic tectum (the number of the animals examined for each group is listed in the panel). Cytoplasmic and yolk injection results are presented for *flt4* and *ccbe1* dgRNPs. (**D**) Experimental workflow of the yolk microinjection experiments for panel (**E**). Injection cocktails containing Cas9 protein with and without the three *vegfab* crRNA:tracrRNA duplexes were injected into the yolk of one-cell stage *Tg(kdrl:EGFP);Tg(kdrl:NLS-mCherry)* embryos. Injected progeny were analyzed for VTA formation at 8 dpf. (**E**) Quantification of ECs at the VTAs within a 5 somite region per animal at 8 dpf after yolk injection of Cas9 protein with and without the three *vegfab* crRNA:tracrRNA duplexes (n=20 for Cas9 controls and n=33 for *vegfab* crRNA1-3). A two-tailed Student’s *t*-test was used to calculate P values. (**F**) Percentage of larvae of indicated treatment with and without VTAs within a 5 somite region per animal at 8 dpf (the number of the animals examined for each group is listed in the panel). Cytoplasmic and yolk injection results are presented for *vegfab* dgRNPs. (**G**) A summary table listing the percentage of larvae of indicated treatment that displayed biallelic mutant vascular phenotypes. A complete absence of FGPs at 6 dpf (*flt4* and *ccbe1*), or of VTAs at 8 dpf (*vegfab*), were defined as biallelic mutant vascular phenotypes. (**H**) A summary table listing the percentage of unviable larvae following the indicated treatment. The number of the animals examined for each treatment is listed in the panel. Fisher’s exact test was used to determine significance between cytoplasmic and yolk injections for *flt4*, *ccbe1*, and *vegfab* in panels (**C**, **F**). However, no statistical difference was detected for any of the gene.

Second, we investigated yolk injections of the three dgRNPs targeting *vegfab* by examining VTA formation at 8 dpf (Figure 5D). We noted that 30 out of 33 larvae (approximately 91%) injected with the three dgRNPs displayed no VTAs. However, the remaining 3 fish showed sprouting vessels at the locations where VTAs normally develop (Figures 5E, 5F). These sprouting vessels were not observed in the larvae injected with the same three *vegfab* dgRNPs via the cytoplasm, suggesting that cytoplasmic injections have a higher efficiency and consistency in this context. Taken together, these results demonstrate that yolk injections of three dgRNPs per gene enabled efficient generation of biallelic F0 knockouts, as recently reported by Kroll et al. [24]. However, our results suggest that cytoplasmic injections perform with even greater efficiency and reproducibility. A summary table listing the percentage of biallelic mutant vascular phenotypes is presented in Figure 5G along with another table summarizing the proportion of unviable larvae observed under each injection condition in Figure 5H.

### Analysis of large genomic deletions and mRNA decay following cytoplasmic dgRNP injections

In order to gain mechanistic insights into how the triple dgRNP injections showed a higher efficiency of biallelic gene disruptions than any of the individual dgRNP injections tested, we investigated genomic deletions and mRNA expression levels following the cytoplasmic injections targeting *flt4* or *vegfab*. To detect genomic deletions flanked by the two farthest dgRNPs, we designed primers upstream and downstream of their respective crRNA1 and crRNA3 target sequences (Figures 6A, 6C). The corresponding dgRNPs containing crRNA1 and crRNA3 were expected to induce approximately 30.4 kb and 1.89 kb genomic deletions in *flt4* and *vegfab* loci, respectively. Upon these expected genomic deletions, the two primers flanking these regions allow for PCR amplification of the expected product size of nearly 400 bp and 430 bp in the deleted loci of *flt4* and *vegfab*, respectively (Figures 6B, 6D). We observed that uninjected siblings and individual dgRNP injections all displayed the PCR products corresponding to their respective WT alleles, but not to their respective genomic deletions (Figures 6E, 6F). In contrast, triple dgRNPs injected samples led to the amplification of the PCR products corresponding to the approximate sizes of the expected deletions (indicated by orange asterisks; Figures 6E–6F’). To determine whether these PCR products indeed resulted from their expected genomic deletions, we sequenced them. We confirmed that they were a result of the expected genomic deletions for both *flt4* and *vegfab*, with slightly different indels integrated into perfect non-homologous end joining of predicted cleavage sites for each injected animal (Figures 6G, 6H). For *flt4*, we detected PCR bands of slightly larger sizes (around 500 bp) in some larval samples (indicated by green asterisks; Figure 6E’), which turned out to be a result of the genomic deletions (approximately 30.3 kb) induced by crRNA2 and crRNA3 (data not shown).

**Figure 6.**
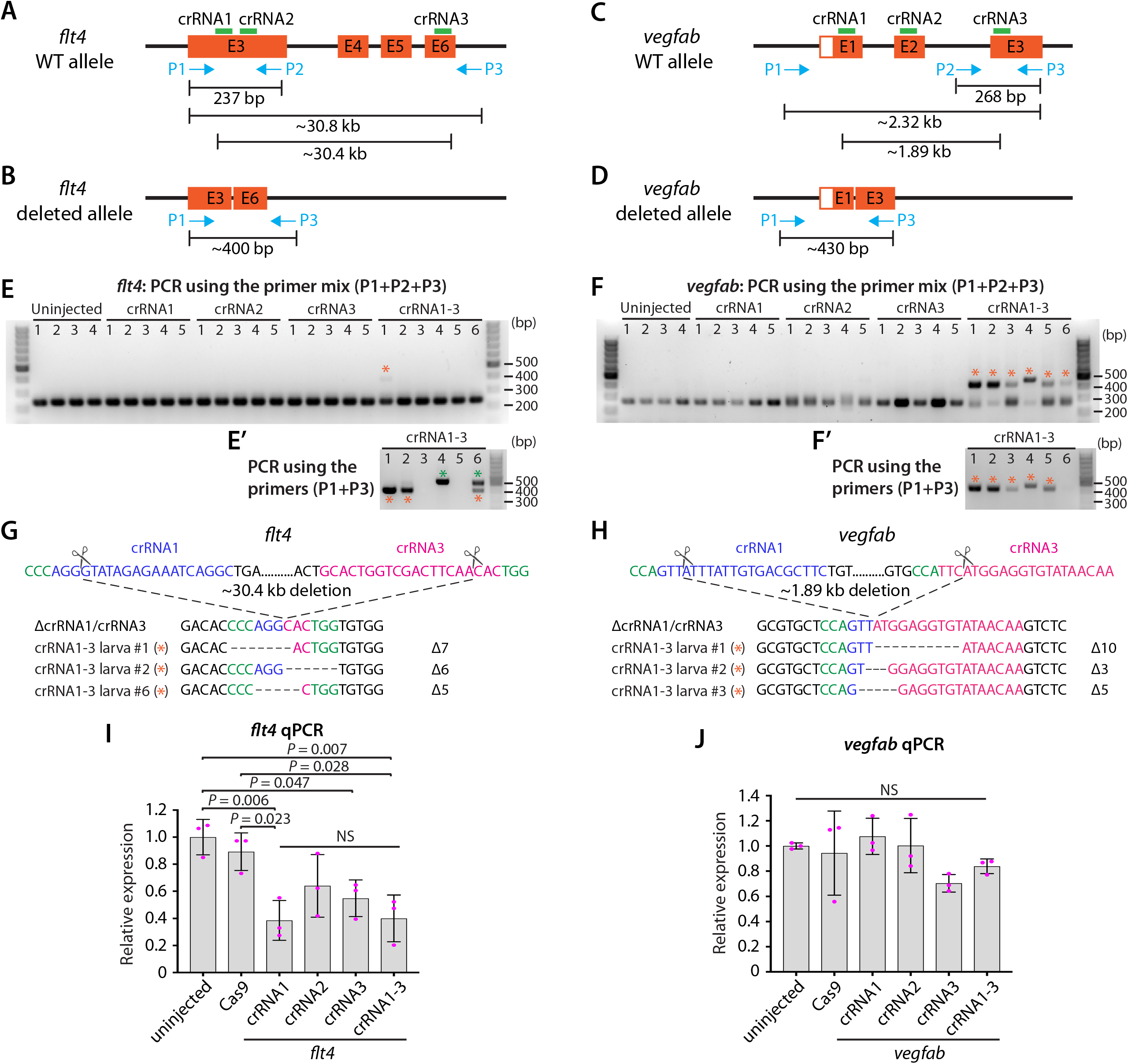
Analysis of large genomic deletions and mRNA decay following cytoplasmic dgRNP injections. (**A**) Diagram of part of the *flt4* genomic locus, indicating approximate locations of the three crRNA target sites and PCR primers. Primer 1 (P1) and primer 2 (P2) were designed to amplify a 237 bp genomic fragment of WT alleles. Primer 3 (P3) was designed far from P1 so these primer pairs cannot efficiently amplify a large genomic fragment (∼30.8 kb) without undergoing genomic deletions (∼30.4 kb) induced by crRNA1 and crRNA3. (**B**) Diagram of the same part of the *flt4* genomic locus after the anticipated genomic deletion (∼30.4 kb) induced by simultaneous targeting of crRNA1 and crRNA3. This genomic deletion is expected to enable PCR amplification of approximately 400 bp of the genomic fragment using primers P1 and P3, which flank these two crRNAs’ target sites. (**C**) Diagram of part of the *vegfab* genomic locus, indicating approximate locations of the three crRNA target sites and PCR primers. Primer 2 (P2) and primer 3 (P3) were designed to amplify a 268 bp genomic fragment of WT alleles. Primer 1 (P1) was designed far from P3 so these primer pairs cannot efficiently amplify a large genomic fragment (∼2.32 kb) without undergoing genomic deletions (∼1.89 kb) induced by crRNA1 and crRNA3. (**D**) Diagram of the same part of the *vegfab* genomic locus after the anticipated genomic deletion (∼1.89 kb) induced by simultaneous targeting of crRNA1 and crRNA3. This genomic deletion is expected to enable PCR amplification of approximately 430 bp of the genomic fragment using primers P1 and P3, which flank these two crRNAs’ target sites. (**E** and **E’**) Genomic DNA from uninjected larvae and those injected with the dgRNPs containing indicated *flt4* crRNAs was amplified with the triple primer mix (P1+P2+P3) (**E**). Uninjected larvae and those injected with individual crRNAs displayed only a 237 bp WT genomic fragment band. However, the WT band and an additional amplicon (asterisks) expected to be generated from genomic deletion alleles were both detected in the genomes of some F0 larvae injected with the triple dgRNPs. The additional amplicons were detected in more efficient manner with PCR using only the primer pairs (P1+P3) (**E’**). The bands marked by orange asterisks were detected at the approximate size of 400 bp, whereas those marked by green asterisks were observed at around 500 bp. These bands were cut out and sequenced. (**F** and **F’**) Genomic DNA from uninjected larvae and those injected with the dgRNPs containing indicated *vegfab* crRNAs was amplified with the triple primer mix (P1+P2+P3) (**F**). Uninjected larvae and those injected with individual crRNAs displayed only a 268 bp WT genomic fragment band. However, the WT band and an additional amplicon (asterisks) expected to be generated from genomic deletion alleles were both detected in the genomes of all F0 larvae injected with the triple dgRNPs. The additional amplicons were detected at the approximate size of 430 bp with PCR using only the primer pairs (P1+P3) (**F’**). These bands were cut out and sequenced. (**G**) Sequence analysis of the PCR products indicated by orange asterisks in **E’** from the triple *flt4* dgRNP-injected larvae. The sequence indicated by ΔcrRNA1/crRNA3 represents expected perfect end joining after simultaneous *flt4* crRNA1 and crRNA3 genetic targeting. The number of base pairs that were apparently deleted from this perfect end joining of predicted cleavage sites is indicated. crRNA1 and crRNA3 target sites are indicated in blue and pink, respectively, along with their PAM sequences (green). (**H**) Sequence analysis of the PCR products indicated by orange asterisks in **F’** from the triple *vegfab* dgRNP-injected larvae. The sequence results from the larvae #1-3 are presented. The sequence indicated by ΔcrRNA1/crRNA3 represents expected perfect end joining after simultaneous *vegfab* crRNA1 and crRNA3 genetic targeting. The number of base pairs that were apparently deleted from to this perfect end joining of predicted cleavage sites is indicated. crRNA1 and crRNA3 target sites are indicated in blue and pink, respectively, along with their PAM sequences (green). (**I**) qPCR analysis of *flt4* mRNA expression levels in 26 hpf embryonic samples of the indicated treatments. Triple dgRNP-injected samples showed significantly reduced *flt4* mRNA levels compared to uninjected and Cas9 injected controls. n=3 biologically independent samples. (**J**) qPCR analysis of *vegfab* mRNA expression levels in 26 hpf embryonic samples of the indicated treatments. No significant difference in *vegfab* mRNA levels was observed across the groups. n=3 biologically independent samples. For qPCR analyses (**I** and **J**), uninjected expression levels were set at 1. A one-way ANOVA followed by Tukey’s post-hoc test was used to calculate P values.

Next, we examined mRNA expression levels by qPCR. We found that *flt4* mRNA levels were reduced to varying degrees at 26 hpf following the triple and any of the single dgRNP injections when compared to uninjected and Cas9 alone-injected control sibling samples (Figure 6I). Injections of dgRNPs containing crRNA1 or all three crRNAs led to prominent reduction (nearly 60%) of *flt4* mRNA compared to uninjected samples (Figure 6I). This qPCR result indicates that triple *flt4* dgRNP injections led to *flt4* mRNA decay, while no significant difference was observed between the triple and any of the single dgRNP injection groups. In the case of *vegfab*, we noted milder mRNA expression differences at 26 hpf. Samples injected with the dgRNPs containing crRNA1 or crRNA2 showed no reduction of *vegfab* mRNA levels compared to those observed in uninjected sibling samples (Figure 6J). Injections of dgRNPs containing crRNA3 or all three crRNAs led to slight reduction (30% and 14%, respectively) of *vegfab* mRNA levels compared to those observed in uninjected samples, although these differences were not statistically significant and were much less prominent than those noted for *flt4*. Moreover, no significant difference in *vegfab* mRNA levels was observed between the triple and any of the single dgRNP injection groups. Collectively, these qPCR results suggest that triple dgRNP injections did not lead to increased mRNA decay compared to single dgRNP injections.

Although no significant difference in *flt4* or *vegfab* mRNA levels after the single and triple dgRNP injections was observed, the HRMA results suggest that individual *flt4* and *vegfab* dgRNPs were highly mutagenic to disrupt their respective target sequences (Figures 2H–J, 4H–J). Furthermore, the combined mutagenic actions of triple *flt4* or *vegfab* dgRNPs led to large genomic deletions (Figures 6E–H). These results together indicate that an increasing number of dgRNPs injected per gene can increase the probability of frameshift mutations via small indels and/or larger genomic deletions, allowing for higher efficiency of biallelic gene disruptions.

### Cytoplasmic injections of two dgRNPs per gene led to variable efficiency of biallelic gene disruptions in F0 animals

Our results showed that cytoplasmic injections of combined three dgRNPs per gene outperformed the induction of biallelic mutant phenotypes (Figure 5G). However, compared to single dgRNP injections, the proportion of unviable larvae observed after triple dgRNP injections was higher (Figure 5H). We next investigated how cytoplasmic injections of two dgRNPs per gene can perform in these two parameters. This investigation allowed us to compare the efficiency of biallelic inactivation as well as unviability between single, dual, and triple dgRNP injections.

To this aim, we focused on *flt4* and *vegfab* and assessed the ability of two dgRNP injections to induce biallelic disruptions using two distinct phenotypic readouts. To make accurate comparisons, we performed dual dgRNP injections in all possible combinations with an equal concentration of Cas9 protein used for single and triple dgRNP injections. We noted that all combinations of dual dgRNP injections targeting *flt4* showed much greater consistency of the “no FGP” phenotype over the optic tectum at 6 dpf than any of the individual dgRNP injections (Figures 7A, 7B, 7E). However, the efficiency of this biallelic mutant phenotype varied among the dgRNP combinations, with the potent crRNA3-containing dgRNP combinations (crRNAs1&3 for 97% and crRNAs2&3 for 100%) outperforming the other dgRNP combination (crRNAs1&2 for 81%) (Figure 7E). The percentage of unviable larvae ranged from 13.6% to 18% (Figure 7F), which is higher than any of the single dgRNP injections, but lower than the triple injections (Figure 5H).

**Figure 7.**
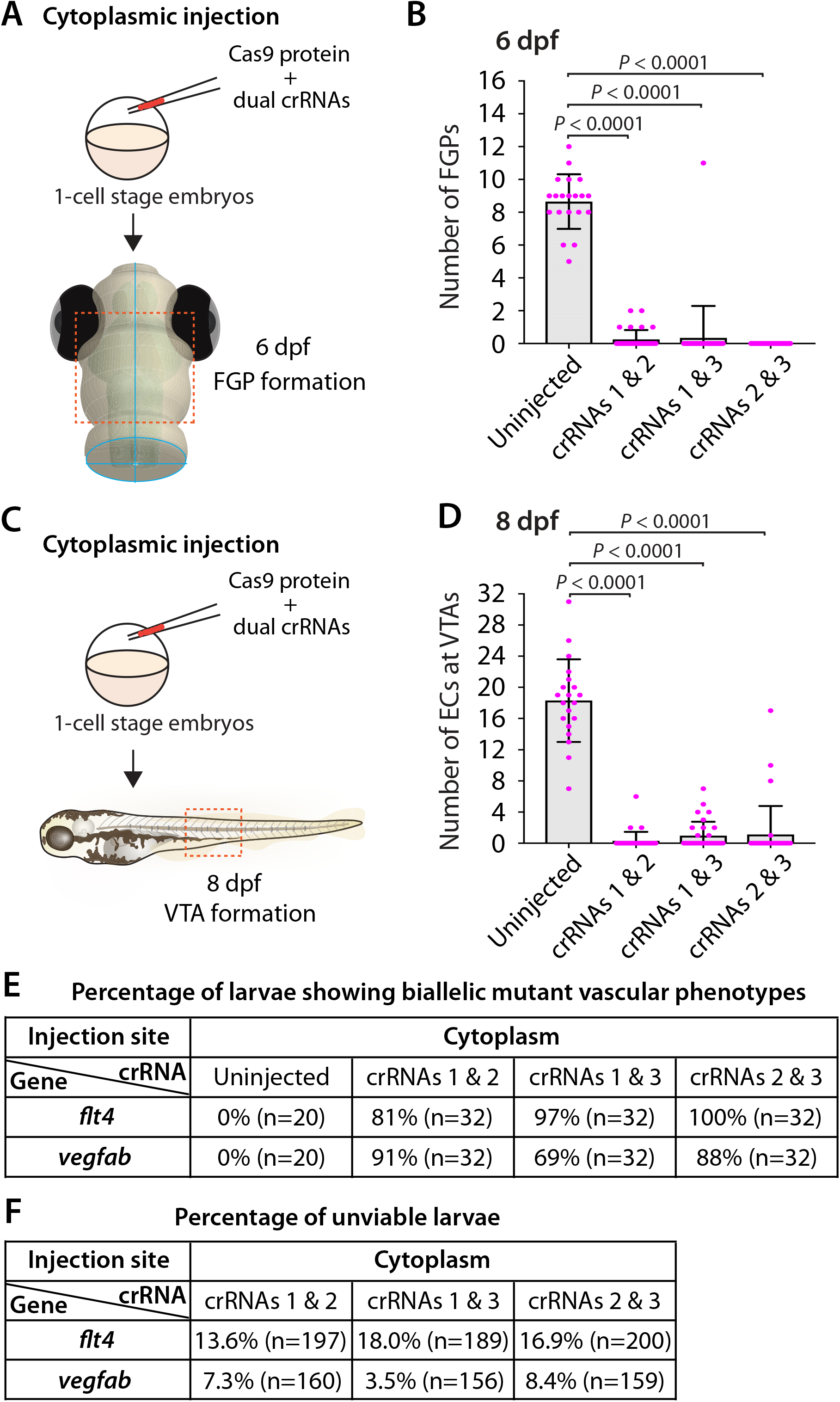
Cytoplasmic injections of two dgRNPs per gene led to variable efficiency of biallelic gene disruptions in F0 animals. (**A**) Experimental workflow of the cytoplasmic microinjection experiments for panel (**B**). Injection cocktails containing Cas9 protein with and without two unique *flt4* crRNA:tracrRNA duplexes were injected into the cytoplasm of one-cell stage *Tg(fli1:nEGFP);Tg(lyve1:DsRed)* embryos. Injected progeny were analyzed at 6 dpf for FGP formation over the optic tectum. (**B**) Quantification of FGPs over the optic tectum at 6 dpf after cytoplasmic injection of Cas9 protein with and without the two indicated *flt4* crRNA:tracrRNA duplexes (n=20 for uninjected controls; and n=32 for *flt4* crRNAs1&2, crRNAs1&3, and crRNAs2&3). Injections of two unique *flt4* dgRNPs led to varying degrees of FGP loss over the optic tectum. (**C**) Experimental workflow of the cytoplasmic microinjection experiments for panel (**D**). Injection cocktails containing Cas9 protein with and without two unique *vegfab* crRNA:tracrRNA duplexes were injected into the cytoplasm of one-cell stage *Tg(kdrl:EGFP);Tg(kdrl:NLS-mCherry)* embryos. Injected progeny were analyzed for VTA formation at 8 dpf. (**D**) Quantification of ECs at the VTAs within a 5 somite region per animal at 8 dpf after yolk injection of Cas9 protein with and without the two indicated *vegfab* crRNA:tracrRNA duplexes (n=20 for uninjected controls; and n=32 for *vegfab* crRNAs1&2, crRNAs1&3, and crRNAs2&3). (**E**) A summary table listing the percentage of larvae of the indicated treatment that displayed biallelic mutant vascular phenotypes. A complete absence of FGPs at 6 dpf (*flt4*), or of VTAs at 8 dpf (*vegfab*), were defined as biallelic mutant vascular phenotypes. (**F**) A summary table listing the percentage of unviable larvae following the indicated treatment. The number of the animals examined for each treatment is listed in the panel. A one-way ANOVA followed by Tukey’s post-hoc test was used to calculate P values for panels (**B**, **D**).

Similarly, we observed that dual dgRNP injections targeting *vegfab* showed overall increased reproducibility of the “no VTA” phenotype at 8 dpf when compared to single dgRNP injections (Figures 5G, 7C, 7D, 7E). However, we noted this phenotype to varying degrees among the dgRNP combinations as observed for the *flt4* experiments. Injections of the potent crRNA2-containing dgRNP combinations (crRNAs1&2 for 91% and crRNAs2&3 for 88%) outperformed the other dgRNP combination (crRNAs1&3 for 69%) (Figure 7E). The proportion of unviable larvae observed after dgRNP injections targeting *vegfab* was low overall (all below 10%), and no drastic differences were noted among the dgRNP injection groups (Figures 5H, 7F). Collectively, these results show that dual dgRNP cytoplasmic injections outperformed single dgRNP injections in the induction of biallelic mutant phenotypes, however, this effect was variable depending on the crRNA combination selected.

### Simultaneous disruptions of functionally redundant genes using a triple dgRNP injection method

Accumulating evidence has indicated that duplicated or related genes in zebrafish play redundant roles in regulating diverse biological processes. This high level of redundancy among functionally-related proteins may prevent one from observing overt phenotypes in single gene knockouts. This phenotypic masking often requires us to generate double and triple genetic mutants to find phenotypes, making phenotypic mutant analyses more laborious. Examining transcriptionally-adapting genes would be informative to narrow down potential candidates that may compensate for the loss of the gene of interest. To quickly test these candidate genes, establishing a rapid and efficient functional platform will be valuable.

We therefore asked whether our mutagenesis approach for generating biallelic F0 knockouts can provide a platform for rapid screening of potential genetic interactions. To test this idea, we examined the genetic interactions between two Vegfr2 zebrafish paralogs, Kdrl and Kdr, in trunk vascular development. Previous studies using morpholinos targeting *kdrl* and *kdr* showed that co-injections of these two morpholinos led to a much more severe defect in aISV formation than their individual injections [31, 39]. A separate study also showed that injecting a *kdr* morpholino into *kdrl* mutants led to a similarly severe aISV formation defect [40]. These studies suggest that Kdrl and Kdr are functionally redundant in regulating aISV formation.

To examine this genetic interaction, we measured aISV lengths in *kdr^+/+^*, *kdr^+/−^*, and *kdr^−/−^* embryos and the same genotypes of embryos in the *kdrl^−/−^* genetic background at 32 hpf (Figure 8A). We noted that while deleting one or two copies of *kdr* in the WT background did not lead to any apparent defect in aISV development (Figures 8B–D, 8H, 8I), doing the same in the *kdrl^−/−^* genetic background dramatically exacerbated aISV formation defects (Figures 8E–G, 8H, 8I). In *kdr^+/+^;kdrl^−/−^* embryos, stalled aISV growth was noted (average aISV lengths 65.8 µm; Figure 8E), consistent with earlier observations (Figures 1B–D, 1H). The same stage of *kdr^+/−^;kdrl^−/−^* embryos displayed more prominent aISV formation defects, leading to much shorter average aISV lengths (16.0 µm) and a partial absence of the dorsal aorta in approximately 9% of fish (Figures 8F, 8H, 8I). Trunk vessel defects were even more severe in *kdr^−/−^;kdrl^−/−^* embryos, resulting in a complete absence of the dorsal aorta in all 16 embryos examined at 32 hpf (Figures 8G, 8H, 8I). This enhanced expressivity of aISV and dorsal aorta phenotypes by deleting one or two copies of *kdr* in the *kdrl^−/−^* genetic background demonstrates the strong genetic interaction between *kdr* and *kdrl*.

**Figure 8.**
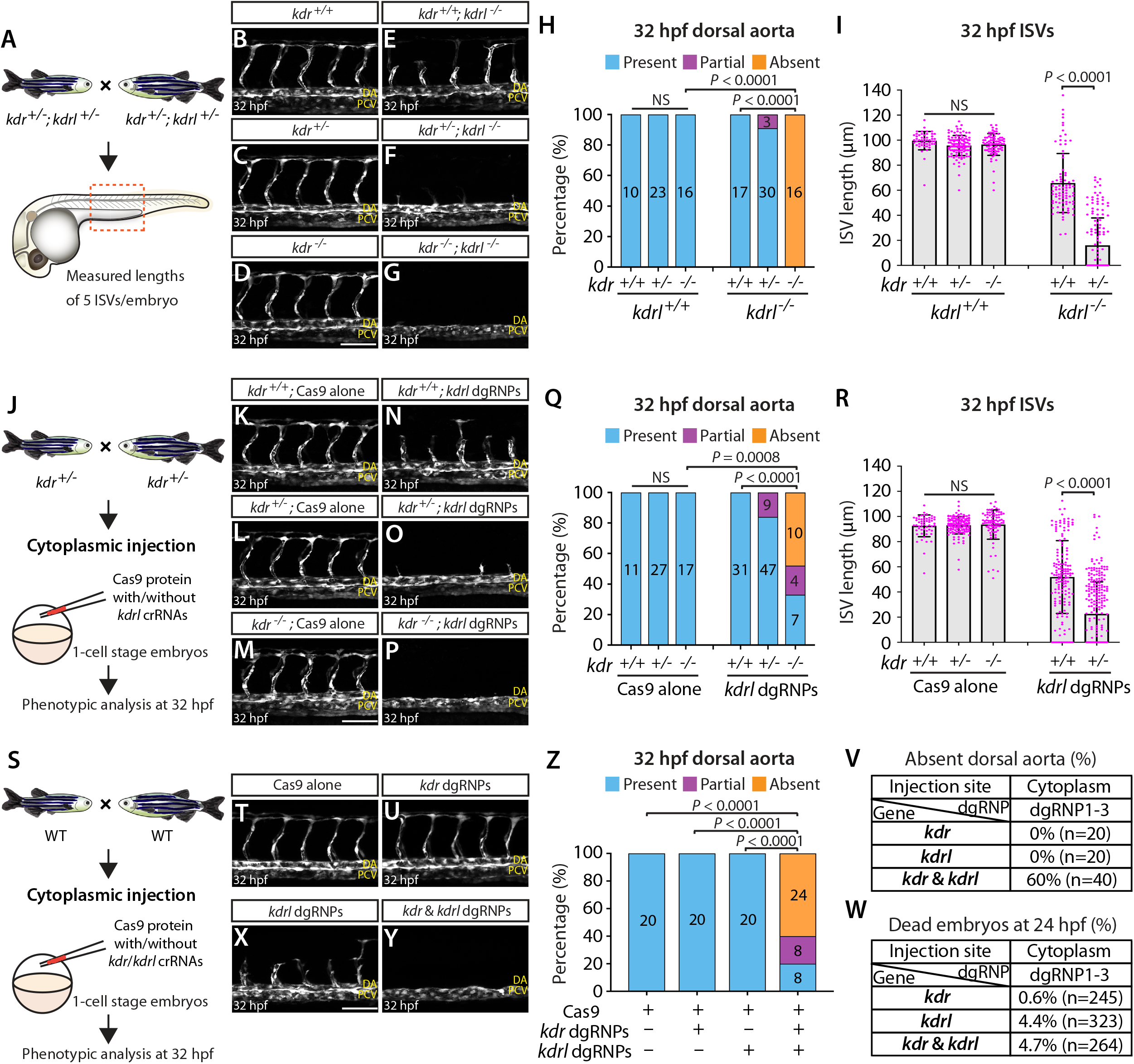
Simultaneous disruptions of the Vegfr2 zebrafish paralog genes using a triple dgRNP injection method. (**A**) Experimental workflow of the phenotypic analysis presented in panels (**B**–**I**). Adult *kdr^+/−^;kdrl^+/−^* fish carrying the *Tg(kdrl:EGFP)* transgene were incrossed, and progeny generated from the crosses were imaged to quantify the formation of the dorsal aorta and aISVs at 32 hpf. (**B**–**G**) Lateral views of *kdr^+/+^* (**B**), *kdr^+/−^* (**C**), *kdr^−/−^* (**D**), *kdr^+/+^;kdrl^−/−^* (**E**), *kdr^+/−^;kdrl^−/−^* (**F**), and *kdr^−/−^;kdrl^−/−^* (**G**) trunk vasculature visualized by *Tg(kdrl:*EGFP) expression at 32 hpf. No obvious trunk vascular defects were observed in *kdr^+/+^*, *kdr^+/−^*, and *kdr^−/−^* embryos (**B**–**D**). *kdr^+/+^;kdrl^−/−^* embryos exhibited stalled aISV growth (**E**), and this aISV growth defect was exacerbated in *kdr^+/−^;kdrl^−/−^* embryos (**F**). In all *kdr^−/−^;kdrl^−/−^* embryos examined at 32 hpf, the dorsal aorta failed to form (**G**). DA: dorsal aorta; PCV: posterior cardinal vein. (**H**) Percentage of 32 hpf embryos of indicated genotype exhibiting the presence, partial presence, and absence of the dorsal aorta (the number of the animals examined per genotype is listed in the panel). (**I**) Quantification of aISV lengths of 32 hpf embryos of the indicated genotypes (n=10 for *kdr^+/+^*, n=23 for *kdr^+/−^*, n=16 for *kdr^−/−^*, n=17 for *kdr^+/+^;kdrl^−/−^*, and n=30 for *kdr^+/−^;kdrl^−/−^*; 5 aISV lengths measured per embryo). *kdr^+/−^;kdrl^−/−^* embryos displayed significantly shorter average aISV lengths than those of *kdr^+/+^;kdrl^−/−^* embryos. (**J**) Experimental workflow of the microinjection experiments for panels (**K**–**R**). Injection cocktails containing Cas9 protein with and without the three *kdrl* crRNA:tracrRNA duplexes were injected into the cytoplasm of one-cell stage embryos generated from incrosses of *kdr^+/−^* fish carrying the *Tg(kdrl:EGFP)* transgene. Injected progeny were imaged to quantify the formation of the dorsal aorta and aISVs at 32 hpf. (**K**–**P**) Lateral trunk views of the 32 hpf *kdr^+/+^* (**K**, **N**), *kdr^+/−^* (**L**, **O**), and *kdr^−/−^* (**M**, **P**) embryos injected with Cas9 protein with (**N**–**P**) or without (**K**–**M**) the three *kdrl* crRNA:tracrRNA duplexes (*kdrl* dgRNPs). Trunk vasculature was visualized by *Tg(kdrl:*EGFP) expression. No obvious trunk vascular defects were observed in *kdr^+/+^*, *kdr^+/−^*, and *kdr^−/−^* embryos injected with Cas9 (**K**–**M**). In contrast, *kdr^+/+^* embryos injected with the triple *kdrl* dgRNPs displayed the stalled aISV phenotype (**N**). Exacerbated aISV defects were observed in the *kdrl* dgRNPs-injected *kdr^+/−^* embryos (**O**). A nearly half of the *kdrl* dgRNPs-injected *kdr^−/−^* embryos exhibited a complete absence of the dorsal aorta (**P**). (**Q**) Percentage of 32 hpf embryos of the indicated genotype and treatment exhibiting the presence, partial presence, and absence of the dorsal aorta (the number of the animals examined per group is listed in the panel). (**R**) Quantification of aISV lengths of 32 hpf embryos of the indicated genotype and treatment (n=11 for Cas9-injected *kdr^+/+^*, n=27 for Cas9-injected *kdr^+/−^*, n=17 for Cas9-injected *kdr^−/−^*, n=31 for *kdrl* dgRNPs-injected *kdr^+/+^*, and n=47 for *kdrl* dgRNPs-injected *kdr^+/−^*; 5 aISV lengths per embryo measured). *kdr^+/−^*embryos injected with the *kdrl* dgRNPs exhibited significantly shorter average aISV lengths than those of the same dgRNPs-injected *kdr^+/+^* fish. (**S**) Experimental workflow of the microinjection experiments for panels (**T**–**W**). Injection cocktails containing Cas9 protein with and without the three *kdr*/*kdrl* crRNA:tracrRNA duplexes were injected into the cytoplasm of one-cell stage embryos generated from incrosses of fish carrying the *Tg(kdrl:EGFP)* transgene. Injected progeny were imaged to observe the formation of the dorsal aorta and aISVs at 32 hpf. (**T**–**Y**) Lateral trunk views of the 32 hpf *Tg(kdrl:EGFP)* embryos injected with Cas9 protein with (**U**–**Y**) or without (**T**) the three *kdr* or *kdrl* crRNA:tracrRNA duplexes (crRNAs), or the combined six *kdr* and *kdrl* crRNAs. No obvious trunk vascular defects were observed in embryos injected with Cas9 (**T**) and the three *kdr* dgRNPs (**U**). In contrast, *kdrl* dgRNPs-injected embryos displayed the stalled aISV phenotype (**X**). The combined injections of the *kdr* and *kdrl* dgRNPs led to a complete absence of the dorsal aorta (**Y**) as observed in 32 hpf *kdr^−/−^;kdrl^−/−^* embryos. (**Z**) Percentage of 32 hpf embryos of the indicated treatment exhibiting the presence, partial presence, and absence of the dorsal aorta (the number of the animals examined per group is listed in the panel). (**V**) Percentage of 32 hpf embryos exhibiting an absence of the dorsal aorta following the cytoplasmic injections of *kdr* or *kdrl* triple dgRNPs, or these combined six dgRNPs (the number of the animals examined per group is listed in the panel). (**W**) Percentage of 24 hpf dead embryos following the cytoplasmic injections of *kdr* or *kdrl* triple dgRNPs, or these combined six dgRNPs (the number of the animals examined per group is listed in the panel). A one-way ANOVA followed by Tukey’s post-hoc test was used to calculate P values for panels (**I**, **R**). Fisher’s exact test was used to calculate P values for panels (**H**, **Q**, **Z**). Scale bars: 50 µm.

Next, we tested this genetic interaction by injecting the three dgRNPs targeting *kdrl* into the cytoplasm of one-cell stage embryos derived from *kdr^+/−^* incrosses (Figure 8J). This approach allowed us to compare aISV lengths in injected embryos of *kdr^+/+^*, *kdr^+/−^*, or *kdr^−/−^* genotypes without generating double mutant animals. We noted that Cas9-injected embryos did not display any overt trunk vessel defects regardless of the *kdr* genotypes (Figures 8K–M, 8Q, 8R). In contrast, simultaneous cytoplasmic injections of the *kdrl* dgRNPs resulted in varying degrees of trunk vessel defects. Subsequent genotyping and quantification results showed that, similar to what we observed in *kdr^+/−^;kdrl^−/−^* embryos, *kdr^+/−^* embryos injected with the three dgRNPs exhibited significantly shorter average aISV lengths (22.6 µm) than those of the dgRNPs-injected *kdr^+/+^* siblings (51.9 µm) (Figures 8N, 8O, 8R). Moreover, approximately 16% of the dgRNPs-injected *kdr^+/−^* embryos displayed a partial loss of the dorsal aorta (Figure 8Q). In the dgRNPs-injected *kdr^−/−^* embryos, we noted that nearly 48% and 19% of them showed a complete and partial loss of the dorsal aorta, respectively (Figures 8P, 8Q). Since an absent dorsal aorta phenotype was observed only in *kdr^−/−^;kdrl^−/−^*, or the dgRNPs-injected *kdr^−/−^*, embryos, this mutagenesis approach allowed efficient biallelic disruption of *kdrl* with this experimental setup as well.

Lastly, we performed cytoplasmic injections of a cocktail containing the three *kdrl* dgRNPs, and three dgRNPs targeting *kdr*, into one-cell stage WT embryos (Figure 8S). This experiment allowed us to test the ability of the triple dgRNP approach in simultaneously targeting two different genes. We first validated the efficacy of each designed crRNA targeting *kdr* by HRMA (Figure S3). Importantly, we verified that all of our designed crRNAs perfectly matched the target sequences within the genome of the corresponding transgenic reporters utilized for quantifications (Figure S4). Next, we doubled the amount of Cas9 protein to make a cocktail containing a total of 6 different dgRNPs. Accordingly, we prepared all control and dgRNP injection cocktails used in this experiment to maintain the equal, increased concentration of Cas9 protein. Under this experimental condition, we observed that all embryos injected with Cas9 alone or with three dgRNPs targeting *kdr* developed the dorsal aorta and aISVs at 32 hpf (Figures 8T, 8U, 8Z), as observed in embryos injected with half the amount of Cas9 protein (Figure 8K), as well as in *kdr^−/−^* embryos (Figure 8D). Injections of the three dgRNPs targeting *kdrl* led to stalled aISV growth at 32 hpf (Figure 8X) while still forming the dorsal aorta (Figures 8X, 8Z), consistent with earlier observations (Figures 1P, 8N). In contrast, embryos injected with a cocktail containing the 6 dgRNPs targeting *kdr* and *kdrl* exhibited severe trunk vascular defects (Figure 8Y), leading to a complete or partial loss of the dorsal aorta in 60% and 20% of the injected embryos, respectively (Figures 8Z, 8V). These results showed that the combined dgRNP injections targeting *kdr* and *kdrl* efficiently inactivated redundant function of these paralog genes without a significant increase in embryonic death (Figure 8W).

Altogether, we provided quantitative evidence that our mutagenesis approach faithfully recapitulated the trunk vascular defects resulting from the genetic interactions between *kdr* and *kdrl*. These results suggest that the cytoplasmic injection approach of three dgRNPs per gene was scalable to target multiple genes, and therefore offer a functional screening platform to test potential genetic redundancy.

## DISCUSSION

Here we present a robust genetic approach that enables a rapid screen of cell autonomous and cell non-autonomous gene function, as well as of genetic redundancy, in F0 zebrafish. This method is user-friendly, since minimal efforts are required for testing the mutagenic activity of designed crRNAs before starting F0 functional screens of the genes of interest (Figure 9). Additionally, these crRNAs can be used to generate stable genetic mutants, including indel and deletion mutants. We were able to successfully generate the *ccbe1* stable mutants shown in this study by utilizing one of the designed crRNAs (Figure S2). Synthetic crRNAs are highly effective, since we designed a total of 17 different synthetic crRNAs in this study, and 16 of them disrupted the corresponding target genomic region as revealed by HRMA. Morpholinos have been utilized to silence gene function in F0 embryos [41, 42], however they cannot be employed to generate genetic mutants. Moreover, the presented method is capable of inactivating gene function over much longer time periods relative to the limited time windows of morpholinos’ gene silencing activity, allowing for wider applications in F0 screens, ranging from early developmental to later physiological and complex behavioral processes. We discuss the advantages of our approach compared to previously reported methods and also describe the potential applications and limitations of this approach.

**Figure 9.**
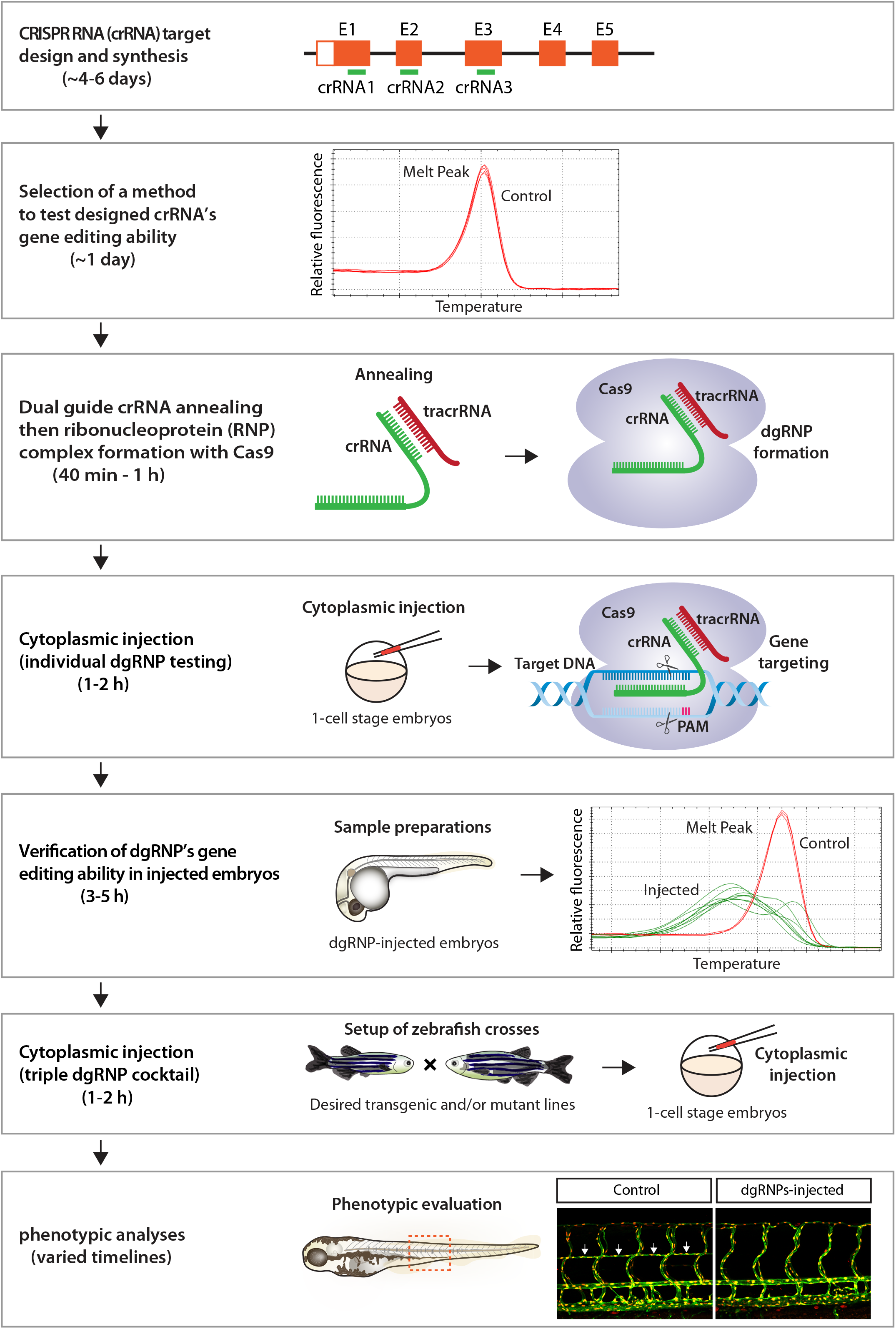
Flowchart summary of triple dgRNP injections for rapid F0 phenotypic screens. This flowchart serves as a quick snapshot of experimental workflow and estimated timelines of the presented method. This method is designed to minimize users’ efforts for crRNA selection, validation, and efficacy testing while maximizing the chance of biallelic disruptions of target genes. Refer to the materials and methods section for detailed methodological descriptions of each step in this flowchart.

### Injections of single, and dual, vs triple dgRNPs per gene

While dgRNP-based mutagenesis has been shown to efficiently disrupt gene function in F0 zebrafish even as single dgRNP injections [23–29], it is becoming clear that anticipated phenotypic detections can depend on target genes and the mutagenic ability of each designed crRNA. This is likely due to varied genetic compensatory responses to mutations in individual loci [12–14, 23] and the variable efficiency of synthetic crRNAs in gene editing [24, 29]. Consistent with these prior reports, we noted that single dgRNP injections often led to variable phenotypes, with only some exhibiting high potency in our experiments (Figure 5G). As such, injection of a single dgRNP per gene can lead to varied biallelic mutagenesis efficiency, resulting in incosistent phenotypic observations in injected F0 animals. Thus, taking a single crRNA injection approach may require significant efforts for selection and validation of crRNAs before starting F0 screens.

A recent study by Kroll et al. reported that an increasing number of injected dgRNPs per gene can achieve higher efficiency of biallelic gene disruptions [24]. They targeted two pigmentation genes, *slc24a5* and *tyr*, and assessed biallelic inactivation caused by yolk injections of 1-4 dgRNPs per gene based on eye pigmentation scoring [24]. They reported that 3 dgRNPs per gene were required for biallelic *slc24a5* disruptions whereas only 2 dgRNPs per gene were needed for *tyr*. However, since the number of the genes and phenotypic readouts they tested were limited, it has left the following isses unresolved: 1) optimal number of dgRNPs per gene in broader biological contexts; 2) choice of injection site; and 3) the ability of such a combined dgRNP injection approach to inactivate diverse gene function (e.g. cell non-autonomous actions).

In this study, we addressed these open questions and present a method that maximizes the efficiency of creating biallelic F0 knockouts, while requiring minimal efforts to check the mutagenic activity of designed crRNAs by HRMA or alternative method(s). To establish this method, we integrated some critical findings from earlier studies, including those made by Hoshijima et al. [23] and Wu et al. [16]. We followed the procedures for dgRNP preparations and injections, including the molar ratio (1:1) and amount (5 fmol) of each injected RNP per embryo, as suggested by Hoshijima et al. [23]. To achieve rigorous phenotypic quantifications, we selected three distinct phenotypic readouts involved in vascular development that are easily quantifiable. Phenotypic comparisons by these quantifiable readouts served to: 1) provide precise information on phenotypic variations between individual animals; 2) allow us to evaluate the ability and efficiency of individual and combined dgRNP injections in inducing knockout phenotypes; and 3) determine the most efficient mutagenesis approach to recapitulate homozygous stable mutant phenotypes.

Our results have added to the recent report by Kroll et al. by providing additional evidence for the broader utility of a three dgRNPs per gene injection approach in abrogating biological processes involving different cell types and gene actions. We determined that cytoplasmic and yolk triple dgRNP injections are highly efficient to induce biallelic gene disruptions with no statistical difference in efficiency between both injection methods. However, our data showed that cytoplasmic injections resulted in slightly greater efficiency and consistency than yolk injections in phenocopying homozygous stable mutant phenotypes. Cytoplasmic dual dgRNPs injections led to enhanced efficiency of inducing biallelic gene disruptions as compared to single dgRNP injections (Figures 5G, 7E). However, we observed the varied effects depending on the targeted genes and dgRNP combination selected, similarly to what is reported by Kroll et al. [24]. A significant difference between their and our triple dgRNP protocols is that we injected nearly half the amount of each dgRNP per embryo (5 fmol) compared to their protocol (9.5 fmol each dgRNP) to achieve strong effects, which will likely help reduce off-target effects and unviable animal rates. Therefore, our presented data provide in-depth quantitative support to establish a further optimized protocol for biallelic gene disruptions in F0 zebrafish.

### A mechanism underlying highly efficient biallelic inactivation after triple dgRNP injections

CRISPR/Cas9-induced DNA double-strand break repair via non-homologous end-joining can result in insertions/deletions of unpredictable length and install a premature termination codon (PTC) [43, 44], which can lead to degradation of the PTC-bearing aberrant mRNA via nonsense-mediated RNA decay [45, 46]. We investigated whether potentially increased probability of frameshift mutations following triple dgRNP injections would result in accelerated mRNA decay. However, our qPCR results for *flt4* and *vegfab* did not show further decreased mRNA levels in samples injected with the respective triple dgRNPs as compared to those following single injections. Although our qPCR experiments were performed only at a single time point, RNA decay alone is less likely to account for the increased efficiency of biallelic disruptions following triple dgRNP injections. However, since expression levels of mRNA and protein are indicated to not always be correlated [47–49], it is possible that functional protein levels were lower in triple dgRNP-injected animals. Despite this possibility, however, increased probability of frameshift mutations appears to be the most likely explanation for the enhanced efficiency of biallelic inactivation following triple dgRNP injections, based on our verification of each designed cRNA for efficient gene targeting and for genomic deletions induced by combined actions of crRNAs. These results support the model proposed by Kroll et al. [24].

### Potential applications and limitations of the presented method

Zebrafish have been used for high-throughput genetic and chemical screenings to identify new genes and pathways fundamental to vertebrate development [50, 51]. The method we present here can be used as an alternative, or supplemental, approach to these genetic and chemical screening methods. For example, it can be used as a secondary genetic assay for validation of key compounds that are identified via phenotype-based small molecule screening. This approach can be also used to quickly test potential candidate genes that have been narrowed down after conducting forward genetic screens and subsequent genetic mapping of causative loci. Thus, our approach can be utilized for a broad range of experimental assays as a primary, secondary, or tertiary method to screen and/or validate the gene of interest, facilitating functional genomic analyses in zebrafish.

Moreover, due to the complex networks of genetic interactions in zebrafish, there is an increasing demand for a rapid screening platform to dissect the roles of duplicated, or adapting, genes with redundant/compensatory functions. As a proof of concept, we provide evidence that our triple dgRNP injection approach addressed the functional redundancy in regulating trunk vascular formation between two duplicated *vegfr2* zebrafish genes, *kdrl* and *kdr*, in a manner comparable to what is observed in their double mutants. Thus, this approach enables rapid functional screening of genetically interacting genes or adapting genes prior to generating zebrafish that harbor multiple mutated genes.

We expect that our approach can also help validate new research tools created in labs, including custom-made antibodies and dominant-negative forms of proteins. For example, the specificity of antibodies can be tested in biallelic F0 knockouts generated via this method if mutants of a gene of interest are not immediately available. In this way, a researcher can forgo at least 3-6 months of testing the specificity of antibodies in mutants that may be imported from elsewhere or generated from scratch. Global or tissue/cell-specific overexpression of dominant-negative forms of proteins has been commonly used to abrogate signaling in zebrafish [52–56]. The efficacy of dominant-negative forms of proteins can be quickly assessed by comparing results to phenotypes observed in biallelic F0 knockouts generated via this method. Hence, this approach will be useful to quickly test new research tools that may need knockout zebrafish samples to address specificity and reliablity.

Potential limitations of this rapid F0 screening approach include off target effects and characterization of genes with unknown phenotypes to date. Cytoplasmic triple dgRNP injections increased the percentage of unviable embryos compared to single dgRNP injections (Figure 5H). However, we did not find significant differences in unviable embryos between cytoplasic and yolk injections. Given the deep sequencing analysis of off target mutations following the triple dgRNP yolk injections reported by Kroll et al. [24], it is unlikely that off target effects will be a serious concern after triple dgRNP cytoplasmic injections, even if sporadic off-target mutations would be induced. One approach to addressing the potential concern of off target effects would be to utilize a different set or combination of crRNAs.

Phenotypic screens of uncharacterized genes may require additional considerations. In our experiments, the average unviability of triple dgRNPs-injected larvae did not exceed 30% for all genes tested, similar to what Kroll et al. reported [24]. Among the viable fish, we decided to analyze only the embryos/larvae that appeared healthy, and we observed a very high induction of biallelic mutant phenotypes. Thus, we anticipate that a vast majority of viable F0 embryos/larvae after injections will exhibit expected biallelic knockout phenotypes. In the case that no expected phenotype are observed, one strategy worth considering would be to generate potential promoter deletion or RNA-less alleles. The successful detection of the *egfl7* mutant vascular phenotype was achieved in this way by Hoshijima et al. [23]. By choosing such a strategy, potential genetic compensation responses that may mask an expected phenotype could be avoided.

In conclusion, our highly efficient mutagenesis protocol for creating biallelic F0 zebrafish knockouts will provide a valuable platform to conduct rapid screening and validation of gene function and redundancy, as well as of research tools, in zebrafish labs. With a careful assessment of off-target effects, a phenotype discovered by triple dgRNP injections is warranted. However, following this F0 screening approach, the results should be recapitulated by generating a stable germline knockout(s).

## Acknowledgements

We thank Drs. Didier Stainier and Nathan Lawson for kindly providing us with fish lines; Don Zeisloft and his team for zebrafish care and husbandry; Drs. Judith Drazba, John Peterson, Gauravi Deshpande, and Ajay Zalavadia for imaging. This work was supported by funding from the National Institutes of Health (R01 NS117510) and start-up funds from the Cleveland Clinic Foundation to R.L.M.

## Author Contributions

R.E.Q. and R.L.M. designed and performed experiments, analyzed data, and wrote the manuscript; L.D.B., S.P., and Z.T. performed experiments and analyzed data. All authors commented on the manuscript.

## Conflict of Interest

The authors declare that they have no conflict of interest.

## Data Availability

All relevant data are provided within the manuscript and its supporting information files.

## Materials and methods

### Zebrafish husbandry and strains

All zebrafish husbandry was performed under standard conditions in accordance with institutional and national ethical and animal welfare guidelines. All zebrafish work was approved by the Cleveland Clinic’s Institutional Animal Care and Use Committee under the protocol number 2018-1970. The following lines were used in this study: *Tg(kdrl:EGFP)^s843^* [57]; *Tg(lyve1:DsRed)^nz101^* [58]; *Tg(fli1:nEGFP)^y7^* [59]; *Tg(kdrl:NLS-mCherry)^is4^* [60]; *kdrl^um19^* [32]; *flt4^um131^* [12]; *vegfab^bns92^* [61]; and *kdr^bns32^* [62]. Adult fish were maintained on a standard 14 h light/10 h dark daily cycle. Fish embryos/larvae were raised at 28.5°C. To prevent skin pigmentation, 0.003% phenylthiourea (PTU) was used beginning at 10-12 hpf for imaging. *flt4* or *ccbe1* dgRNP-injected larvae analyzed at 10 dpf were transferred to a tank containing approximately 250 mL system water supplemented with 0.003% PTU (up to 25 larvae/tank) and fed with Larval AP100 (<50 microns dry diet, Zeigler) starting at 5 dpf. *vegfab* dgRNP-injected larvae analyzed at 8 and 12 dpf were transferred to a tank containing approximately 250 mL system water without PTU (up to 25 larvae/tank) and fed with Larval AP100 (<50 microns dry diet, Zeigler) starting at 5 dpf.

### Genotyping of mutants

Genotyping of *kdrl^um19^*, *flt4^um131^*, *vegfab^bns92^*, and *kdr^bns32^* mutant fish was performed by high-resolution melt analysis of PCR products using the following primers:

*kdrl um19* forward: 5’ – TGCTTCCTGATGGAGATACACACC – 3’

*kdrl um19* reverse 5’ – TGCAAATGAGTGTGAGTGTCCCAC – 3’

*flt4 um131* forward: 5’ – GACCATCTTCATAACAGACTCTG – 3’

*flt4 um131* reverse: 5’ – GGATCTGAAACCAGACATGGTAC – 3’

*vegfab bns92* forward: 5’– GTGCTGGGTGCTGCAATG – 3’

*vegfab bns92* reverse: 5’– CCAAGGTAATGTTGTATGTGACG – 3’

*kdr bns32* forward: 5’ – GACCTCACCCTGAGTCCACA – 3’

*kdr bns32* reverse 5’ – GCGGTGCAGTTGAGTATGAG – 3’

### Generation and genotyping of *ccbe1* mutants

The *ccbe1^lri97^* mutant allele was generated by targeted genome editing using the CRISPR/Cas9 system as follows: We targeted the first exon of *ccbe1*, which encodes the N-terminal part of the Ccbe1 protein-coding sequences prior to the EGF domain. The *ccbe1^lri97^* mutant allele harbors a 6 base pair deletion and a 1 base pair insertion. The *ccbe1^lri97^* allele is predicted to carry a frameshift mutation that alters the reading frame of Ccbe1 at threonine residue 26, leading to a premature stop codon after 40 missense amino acids. The following sequences of *ccbe1* CRISPR RNA (crRNA) were used to generate the mutants:

*ccbe1* crRNA: TTCTCCTCTCGGAAAGTCCA

Ribonucleoprotein (RNP) complexes of *ccbe1* crRNA, trans-activating RNA (tracrRNA), and Cas9 protein were prepared as described in detail in the method section of crRNA preparations and injections. Injection cocktails (4 μL) containing the crRNA:tracrRNA:Cas9 RNP complexes were prepared as follows: 0.4 μL 25 μM crRNA:tracrRNA, 0.4 μL 25 μM Cas9 protein stock, 2.8 μL H_2_O, and 0.4 μL 0.5% phenol red solution (Sigma). Prior to microinjection, injection cocktails were incubated at 37°C for 5 min and then kept at room temperature. Approximately 1 nL of the injection cocktails was injected into the cytoplasm of one-cell stage embryos. Genotyping of *ccbe1^lri97^* mutant fish was performed by HRMA of PCR products using the following primers:

*ccbe1 lri97* forward: 5’ – ACTGTTCTCATCGGGAGCTCCGTG – 3’

*ccbe1 lri97* reverse: 5’ – AGCACTTTACCTGTCTACATCCTC – 3’

### High-resolution melt analysis (HRMA)

Embryos injected with individual crRNA-containing RNP complexes were subjected to genomic DNA extraction at 24 hpf. Primer sequences used to analyze the efficacy of individual crRNAs in their injected embryos are listed in Table 1. Fish embryos/larvae carrying stable genetic mutations were collected and subjected to genomic DNA extraction immediately after confocal imaging. A CFX96 Touch Real-Time PCR Detection System (Bio-Rad) was used for PCR reactions and HRMA. Precision Melt Supermix for high-resolution melt analysis (Bio-Rad) was used in these experiments. PCR reaction protocols were: 95°C for 2 min, 46 cycles of 95°C for 10 s, and 60°C for 30 s. Following the PCR, a high-resolution melt curve was generated by collecting EvaGreen fluorescence data in the 65–95°C range. The analyses were performed on normalized derivative plots.

**Table 1.**
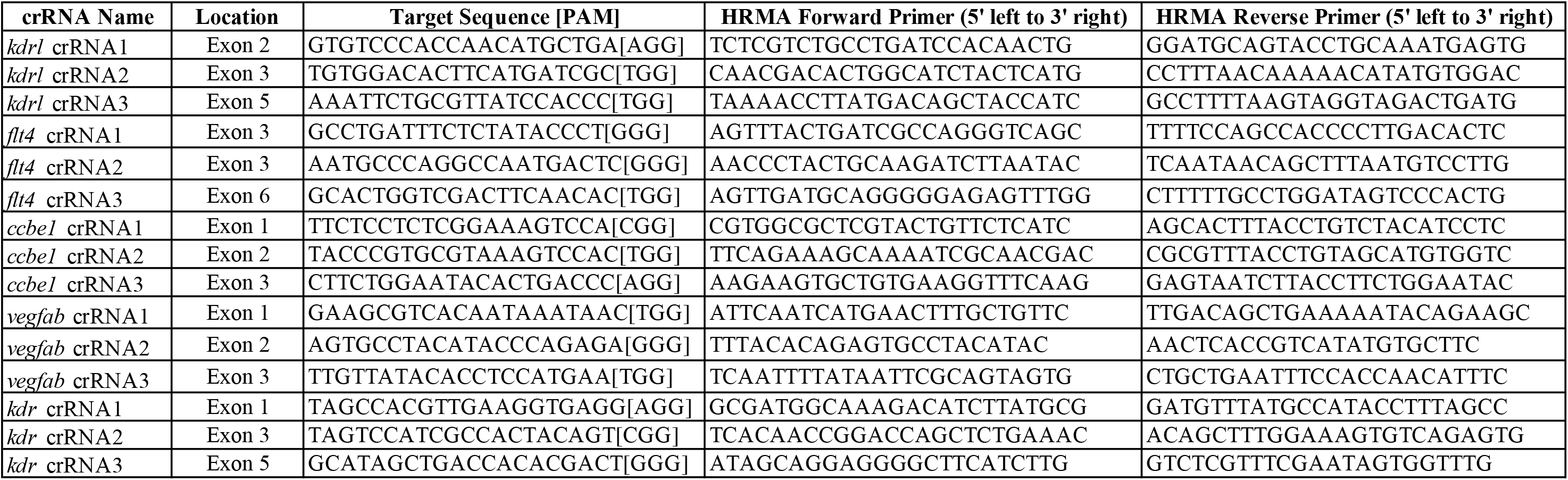
crRNA target sites, genomic locations, and HRMA primers used to analyze crRNA-injected embryos.

### crRNA target site design

crRNA target sites were selected using the following web programs: Predesigned (https://www.idtdna.com/site/order/designtool/index/CRISPR_PREDESIGN) and Custom (https://www.idtdna.com/site/order/designtool/index/CRISPR_CUSTOM) Alt-R® CRISPR-Cas9 guide RNA design that is operated by Integrated DNA Technologies, IDT. These web programs rank potential crRNA target sites based on predicted on-target and off-target scores. crRNA target sites for each gene were selected based on this prediction, with higher priority given to those with less off-target effects. All crRNA target sites are listed in Table 1. Only adults carrying verified target sequences (Figure S4) were used for gene editing experiments and for quantifications.

### crRNA preparations and injections

Designed crRNAs and tracrRNA were all purchased from IDT. dgRNP complexes composed of crRNA, tracrRNA, and Cas9 protein were prepared as follows: Duplex buffer (IDT) was used to make a 100 μM stock solution of individual crRNA and tracrRNA. To generate crRNA:tracrRNA duplex complexes (50 μM), each crRNA stock solution (100 μM) was mixed with an equal volume of tracrRNA (100 μM) and annealed in a thermal cycler: 95°C for 5 min; cooling at 0.1°C/sec to 25°C; 25°C for 5 min; 4°C hold. 50 μM crRNA:tracrRNA duplex solution was diluted in duplex buffer to generate 25 μM stock solution, which was stored at −20°C. Target genomic sequences and locations of each designed crRNA are listed in Table 1.

For experiments shown in Figure 8S–W, a 50 μM stock solution of Cas9 protein (S.p. Cas9 nuclease, V3, IDT) was prepared by diluting a 62 μM original Cas9 protein sample from IDT with Cas9 buffer (20 mM HEPES-NaOH (pH 7.5), 350 mM KCl, and 20% glycerol). Diluted Cas9 protein stocks were aliquoted and stored at −80°C. Injection cocktails (4 μL) containing the three different *kdr* and *kdrl* dgRNP complexes were prepared as follows: 0.4 μL 25 μM *kdr* crRNA1:tracrRNA, 0.4 μL 25 μM *kdr* crRNA2:tracrRNA, 0.4 μL 25 μM *kdr* crRNA3:tracrRNA, 0.4 μL 25 μM *kdrl* crRNA1:tracrRNA, 0.4 μL 25 μM *kdrl* crRNA2:tracrRNA, 0.4 μL 25 μM *kdrl* crRNA3:tracrRNA, 1.2 μL 50 μM Cas9 protein stock, and 0.4 μL 0.5% phenol red solution. Injection cocktails (4 μL) containing three different *kdr* or *kdrl* dgRNP complexes were prepared as follows: 0.4 μL 25 μM crRNA1:tracrRNA, 0.4 μL 25 μM crRNA2:tracrRNA, 0.4 μL 25 μM crRNA3:tracrRNA, 1.2 μL 50 μM Cas9 protein stock, 1.2 μL H_2_O, and 0.4 μL 0.5% phenol red solution. Control injection cocktails (4 μL) without cRNA:tracrRNA duplexes were prepared as follows: 1.2 μL 50 μM Cas9 protein stock, 2.4 μL H_2_O, and 0.4 μL 0.5% phenol red solution. All injection cocktails contained an equal concentration of Cas9 protein to allow for direct quantitative comparisons between these injection groups.

For the remaining dgRNP injection experiments, a 25 μM stock solution of Cas9 protein was prepared by diluting a 62 μM original Cas9 protein sample from IDT with the same Cas9 buffer described earlier. Diluted Cas9 protein stocks were aliquoted and stored at −80°C. Injection cocktails (4 μL) containing three different dgRNP complexes were prepared as follows: 0.4 μL 25 μM crRNA1:tracrRNA, 0.4 μL 25 μM crRNA2:tracrRNA, 0.4 μL 25 μM crRNA3:tracrRNA, 1.2 μL 25 μM Cas9 protein stock, 1.2 μL H_2_O, and 0.4 μL 0.5% phenol red solution. Injection cocktails (4 μL) containing two different dgRNP complexes were prepared as follows: 0.4 μL 25 μM first crRNA:tracrRNA, 0.4 μL 25 μM second crRNA:tracrRNA, 1.2 μL 25 μM Cas9 protein stock, 1.6 μL H_2_O, and 0.4 μL 0.5% phenol red solution. Injection cocktails (4 μL) containing individual dgRNP complexes were prepared as follows: 0.4 μL 25 μM crRNA:tracrRNA, 1.2 μL 25 μM Cas9 protein stock, 2 μL H_2_O, and 0.4 μL 0.5% phenol red solution. Control injection cocktails (4 μL) without cRNA:tracrRNA duplexes were prepared as follows: 1.2 μL 25 μM Cas9 protein stock, 2.4 μL H_2_O, and 0.4 μL 0.5% phenol red solution. All injection cocktails contained an equal concentration of Cas9 protein to allow for direct quantitative comparisons between single, dual, and triple dgRNP injections.

For all injection experiments, injection cocktails were freshly prepared on the day of injection and were incubated at 37°C for 5 min and kept at room temperature prior to microinjection. Approximately 2 nL of the injection cocktails was injected into the cytoplasm or yolk of one-cell stage embryos. For phenotypic quantifications throughout this study, we chose to analyze only the embryos/larvae that did not display obvious gross morphological defects regardless of being injected or uninjected. Phenotypic quantification results represent data of the independent biological samples collected from injections performed on different days.

### Detection and evaluation of genomic deletion mutations

Genomic DNA was extracted from individual larvae at 3 dpf. Each larva was collected in a PCR tube, anesthetized, and incubated in 25 μL base solution (25 mM KOH and 200 μM EDTA) at 95°C for 30 min. After cooling down to room temperature, 25 μL of neutralization solution (40 mM Tris-HCl) was added. Genomic deletions that brought *flt4* or *vegfab* crRNA1 and crRNA3 target sites close together were detected following PCR amplification with primers that flanked those sites. Primer sequences are listed in Table S2. The PCR reaction mixture was prepared for a 10 μL reaction volume using a GoTaq® DNA Polymerase PCR mix (Promega): 2 μL 5X Green GoTaq® Reaction Buffer, 0.5 μL dNTP mix (2.5 mM each), 0.5 μL three primer mix (P1, P2, and P3: 10 μM each), 0.1 μL GoTaq® DNA Polymerase, 5.9 μL H_2_O, and 1 μL 1:20 diluted genomic DNA in H_2_O. The PCR program was: 95°C for 3 min; then 35 cycles of: 95°C for 30 s, 60°C for 30 s, 72°C for 30 s; then 72°C for 5 min; 4°C hold. To determine genomic deletion mutations, the PCR reaction mixture was assembled in the same way described earlier, except the use of two primer mix (P1 and P3: 10 μM each) instead of the three primer mix, and subjected to the PCR program: 95°C for 3 min; then 40 cycles of: 95°C for 30 s, 60°C for 30 s, 72°C for 30 s; then 72°C for 5 min; 4°C hold. After PCR, bands of the approximate size expected to be generated from genomic deletion alleles were cut from agarose gels, purified using a GeneJET Gel Extraction Kit (Thermo Fisher Scientific, Cat. No. K0692), and sequenced via tube sequencing services (Eurofins Genomics). The sequence chromatograms and results were analyzed using SnapGene and ApE software.

### Verification of crRNA target sites and PAM sequences in the genomes of zebrafish transgenic lines

Genomic DNA was extracted from amputated caudal fins of individual adult zebrafish that carried *Tg(kdrl:EGFP)^s843^;Tg(kdrl:NLS-mCherry)^is4^* or *Tg(fli1:nEGFP)^y7^;Tg(lyve1:DsRed)^nz101^* reporters. The caudal fins amputated from 3 mating pairs of adults carrying either reporter were subjected to genomic DNA extraction. Genome sequences containing crRNA target sites and PAM sequences were amplified with primer pairs listed in Table 2. The PCR reaction mixture was prepared for a 10 μL reaction volume using a GoTaq® DNA Polymerase PCR mix (Promega): 2 μL 5X Green GoTaq® Reaction Buffer, 0.5 μL dNTP mix (2.5 mM each), 0.5 μL forward and reverse primer mix (10 μM each), 0.1 μL GoTaq® DNA Polymerase, 5.9 μL H_2_O, and 1 μL 1:20 diluted genomic DNA in H_2_O. The PCR program was: 95°C for 3 min; then 40 cycles of: 95°C for 30 s, 60°C for 30 s, 72°C for 30 s; then 72°C for 5 min; 4°C hold. After PCR, single bands of the expected size were cut from agarose gels, purified using a GeneJET Gel Extraction Kit (Thermo Fisher Scientific, Cat. No. K0692), and sequenced via tube sequencing services (Eurofins Genomics). The sequence chromatograms and results were analyzed using SnapGene and ApE software.

**Table 2.**
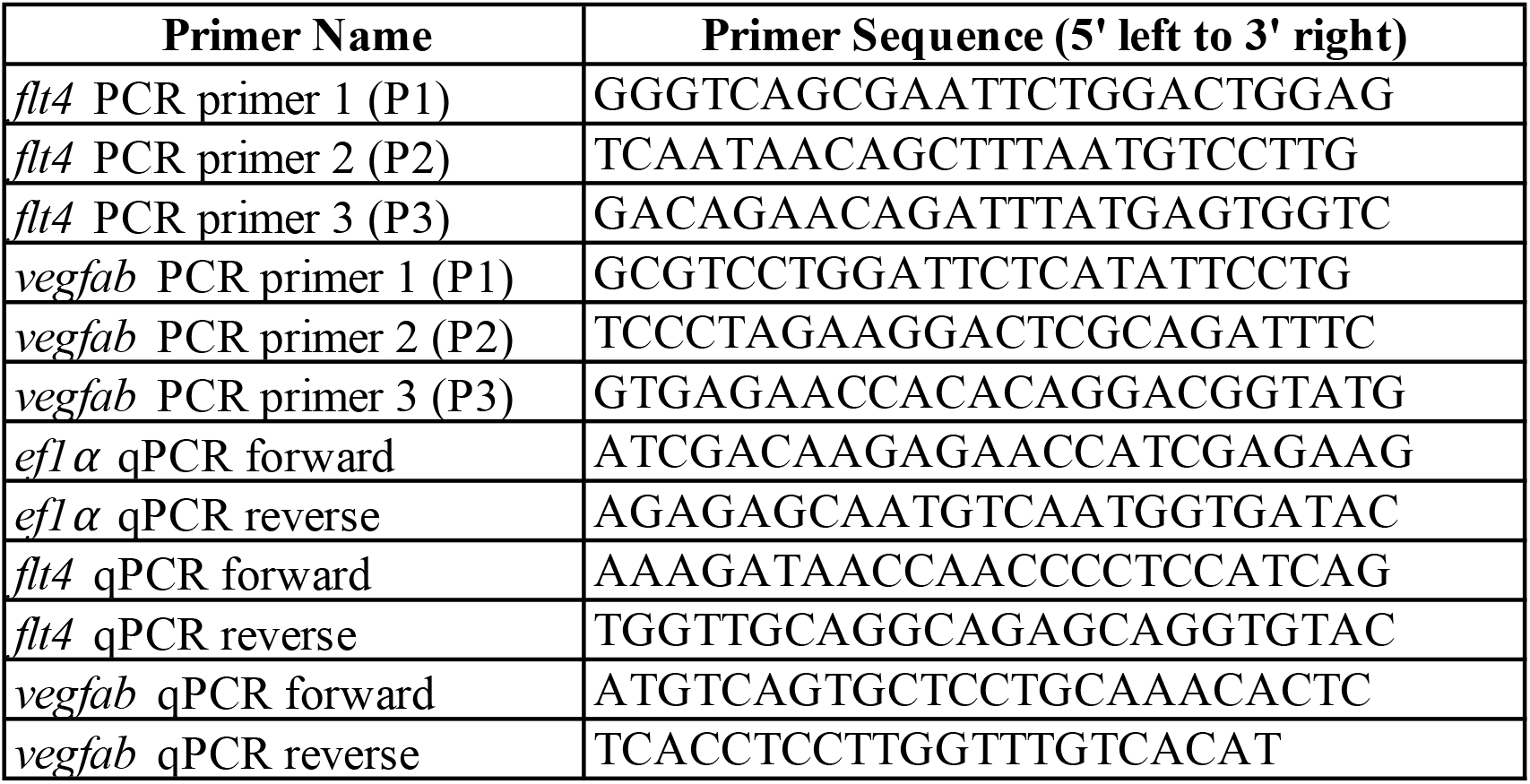
PCR and qPCR primers used to examine genomic deletions and mRNA expression.

**Table 3.**
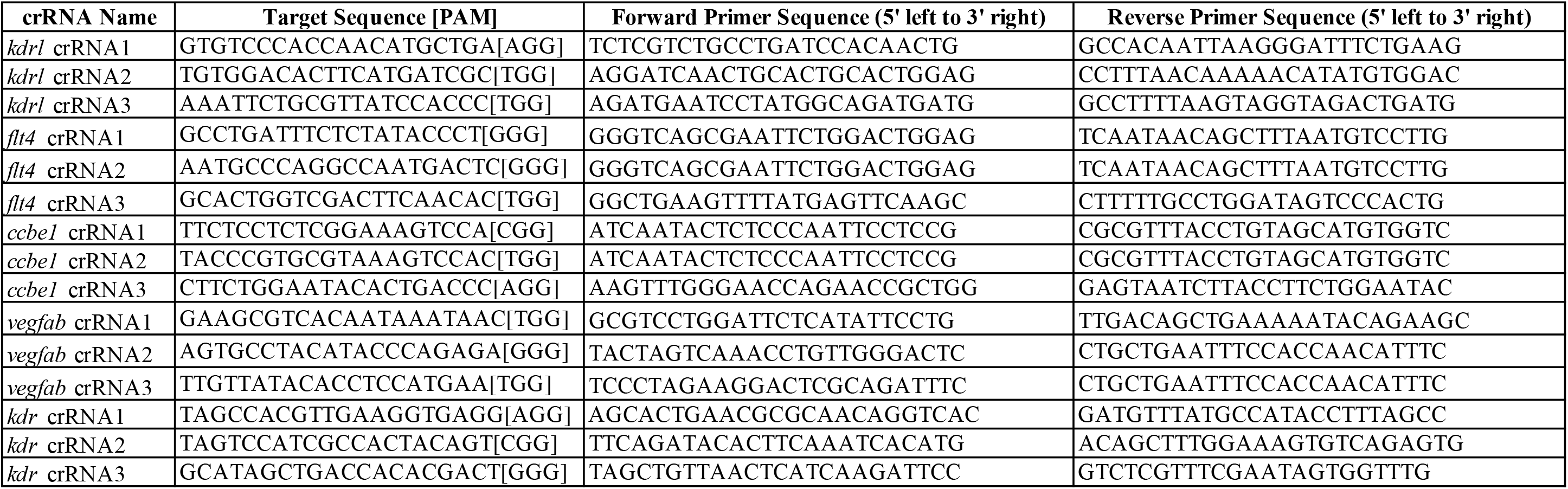
PCR primer pairs used to amplify genomic fragments containing crRNA target sequences within the genome of zebrafish strains.

### qPCR expression analysis

qPCR was performed on cDNA obtained from 26 hpf whole embryos after injections at the 1-cell stage with individual or triple crRNA-containing dgRNP complexes targeting *flt4* or *vegfab*. For each uninjected and injected treatment sample, 5 embryos were pooled for RNA isolation. Embryos were homogenized in QIAzol Lysis Reagent and subjected to RNA isolation using miRNeasy Micro Kit (Qiagen, Cat. No. 217084). Isolated RNA was then purified using RNA Clean & Concentrator-5 kit (Zymo Research, Cat. No. R1013). 500 ng RNA was used for cDNA synthesis for each sample using the iScript cDNA Synthesis kit (Bio-Rad, Cat. No. 1708890). A CFX96 Touch Real-Time PCR Detection System (Bio-Rad) was used for qPCR experiments, with an input of 10 ng cDNA per reaction. Gene expression levels were normalized relative to that of the reference gene *ef1α*. All reactions were performed in technical duplicates, and the results represent biological triplicates. Fold changes were calculated using the 2^−ΔΔCt^ method. Primer sequences used for the qPCR experiments are listed in Table 2.

### Confocal and stereo microscopy

A Leica TCS SP8 confocal laser scanning microscope (Leica) was used for live imaging. Live zebrafish embryos and larvae were anaesthetized with a low dose of tricaine, embedded in a layer of 1% low melt agarose in a glass-bottom Petri dish (MatTek), and imaged using a 25X water immersion objective lens. Leica Application Suite X (LAS X) software (Version 3.7.0.20979) was used for image acquisition and analysis. An SMZ-18 stereomicroscope (Nikon) was used for brightfield images of anaesthetized fish. NIS-Elements BR Imaging Software (Version 5.10.01) was used for image acquisition and analysis.

### Quantification of arterial intersegmental vessel (aISV) lengths and dorsal aorta formation

Fish embryos carrying the *Tg(kdrl:EGFP)^s843^* transgene were used to quantify the lengths of aISVs and dorsal aorta formation at 32 hpf. For the quantification of aISVs in *kdrl^+/+^*, *kdrl^+/−^*, and *kdrl^−/−^* embryos, *kdrl^um19^* heterozygous adults carrying the *Tg(kdrl:EGFP)* reporter were incrossed. For the genetic interaction experiments, *kdrl^um19/+^;kdr^bns32/+^* adults carrying the *Tg(kdrl:EGFP)* reporter were incrossed. For the *kdrl* dgRNP injection experiments in Figures 8J–R, WT or *kdr^bns32^* heterozygous adults carrying the *Tg(kdrl:EGFP)* reporter were incrossed, and injection cocktails containing Cas9 protein with and without three *kdrl* crRNA(s):tracrRNA duplexes were injected into the cytoplasm of one-cell stage progeny generated from these crosses. For the *kdr* and *kdrl* dgRNP injection experiments in Figures 8S–W, WT adults carrying the *Tg(kdrl:EGFP)* reporter were incrossed, and injection cocktails containing Cas9 protein with and without three *kdr*/*kdrl* crRNA(s):tracrRNA duplexes were injected into the cytoplasm of one-cell stage progeny generated from these crosses. For each fish, the lengths of 5 aISVs in the left side of the trunk region directly anterior to the anal opening were measured using the scale bar tool in LAS X software. aISV measurements began at the dorsal surface of the dorsal aorta, ending at either the dorsal tip of the aISV or at the connection of the aISV and dorsal longitudinal anastomotic vessel. The presence, partial presence, and absence of the dorsal aorta was examined in the same trunk region used for aISV length measurements.

### Quantification of perivascular FGPs in the dorsal meningeal compartment over the optic tectum

*Tg(fli1:nEGFP)^y7^;Tg(lyve1:DsRed)^nz101^* reporter fish were used to quantify the number of FGPs in the dorsal meningeal compartment over the optic tectum at 6 dpf. For the quantification of FGPs in *flt4^+/+^*, *flt4^+/−^*, and *flt4^−/−^* larvae, *flt4^um131^* heterozygous adults carrying the *Tg(fli1:nEGFP);Tg(lyve1:DsRed)* reporters were incrossed. For the quantification of FGPs in *ccbe1^+/+^*, *ccbe1^+/−^*, and *ccbe1^−/−^* larvae, *ccbe1^lri97^* heterozygous adults carrying the *Tg(fli1:nEGFP);Tg(lyve1:DsRed)* reporters were incrossed. For the *flt4* and *ccbe1* dgRNP injection experiments, *Tg(fli1:nEGFP);Tg(lyve1:DsRed)* adults were incrossed, and injection cocktails containing Cas9 protein with and without crRNA(s):tracrRNA duplex complexes were injected into the cytoplasm or yolk of one-cell stage progeny generated from these crosses as indicated in related figures and legends. *Tg(fli1:*nEGFP*);Tg(lyve1:*DsRed)-double positive FGPs were quantified by counting the number of *Tg(fli1:*nEGFP)^+^ nuclei in these double-positive cells. LAS X software was utilized to define the region used for this quantification. In the maximum projection confocal images, 150 μm distances from the midline on both sides of the head were measured to set the defined caudolateral area for quantifications across samples (see Figure S1 for the defined areas used to quantify FGPs in this study). The midline corresponds to the region at which the bilateral mesencephalic veins converge. The most lateral point of this line denoted the cutoff point for countable double-positive cells; cells medial to this line were included in the count. Next, 75 μm distances from the midline on both sides were measured to set the defined rostrolateral area for quantifications. The rostral extension of the *Tg(lyve1:*DsRed*)^+^* loop intersects this line, and this point serves as the most lateral cutoff point for quantification of double positive cells; cells medial to this line were included in the count. Refer to Figure S1 for further information.

### Quantification of the number of the endothelial cells that form vertebral arteries (VTAs)

*Tg(kdrl:EGFP)^s843^;Tg(kdrl:NLS-mCherry)^is4^* reporter fish were used to quantify the number of endothelial cells (ECs) that form VTAs at 8 dpf. For the quantification of EC number in *vegfab^+/+^*, *vegfab^+/−^*, and *vegfab^−/−^* larvae, *vegfab^bns92^* heterozygous adults carrying the *Tg(kdrl:EGFP);Tg(kdrl:NLS-mCherry)* reporters were incrossed. For the *vegfab* dgRNP injection experiments, *Tg(kdrl:EGFP);Tg(kdrl:NLS-mCherry)* adults were incrossed, and injection cocktails containing Cas9 protein with and without crRNA(s):tracrRNA duplex complexes were injected into the cytoplasm or yolk of one-cell stage progeny generated from these crosses as indicated in related figures and legends. *Tg(kdrl:*EGFP*);Tg(kdrl:*NLS-mCherry)-double positive ECs that comprise VTAs were quantified by counting the number of *Tg(kdrl:*NLS-mCherry)^+^ nuclei in these double-positive ECs. The 5 somites directly anterior to the anal opening were examined on both sides of the body (defined as a 5 somite region) to quantify the number of ECs comprising the VTAs at 8 dpf.

### Quantification of unviable embryos/larvae

For experiments shown in Figure 8S–W, the numbers of live and dead embryos were recorded at 24 hpf, and the percentage of dead embryos was calculated for each injection group. Uninjected sibling embryos from the breeding pairs utilized for each injection were used to determine the percentage of unfertilized eggs in individual clutches at 24 hpf. This percentage was subtracted from the percentage of dead embryos in each injection group to determine only the effects that were induced by injections. When this subtraction resulted in negative numbers, we considered this death rate as zero percent since this result indicated that no adverse effects on viability were induced by injections. The average death percentage of the sibling embryos injected with Cas9 alone (10 fmol) for these experiments was 3.0%.

For the remaining dgRNP injection experiments that counted unviable embryos/larvae, the percentage of unviable embryos/larvae was calculated based on the total number of dead and dysmorphic larvae (exhibiting developmental deficits that were not associated with the anticipated phenotypes) after 24 hpf. Dead embryos at 0 dpf were excluded since they were likely either unfertilized eggs or were damaged by the injection needle during microinjection procedures. At 72 hpf, the numbers of apparently healthy-looking and dead/dysmorphic (deformed, curved and/or short body, and severe peri-cardiac edema) embryos were recorded. Since *flt4* and *ccbe1* gene disruptions could lead to mild peri-cardiac edema (potential phenotype), embryos with such mild edema were considered not to be dysmorphic. After counting at 72 hpf, dead/dysmorphic numbers were divided by the total number of live embryos at 24 hpf to obtain a percentage of unviable embryos. The same method described earlier was used to calculate unviability. The average unviability percentage of the sibling embryos injected with Cas9 alone (5 fmol) for these experiments was 4.4%.

For all injection experiments, at least four clutches of 1-cell stage embryos were injected, and then subjected to quantification. The number of embryos per clutch ranged from 20 to 80, with most clutches containing 40-60 embryos. Dead and unviable embryo percentages calculated from each clutch were averaged and presented along with the total number of animals examined in Figures 5H, 7F, and 8W.

### Statistical analysis

Statistical differences for mean values among multiple groups were determined using a one-way analysis of variance (ANOVA) followed by Tukey’s or Dunnett’s multiple comparison test. Statistical differences for mean values between two groups were determined using a two-tailed Student’s *t*-test. Fisher’s exact test was used to determine significance when comparing the degree of penetrance of observed phenotypes. Statistical analyses were performed using GraphPad Prism 8.1.1. The criterion for statistical significance was set at *P* < 0.05. Error bars are SD.

---

**Figure S1.**
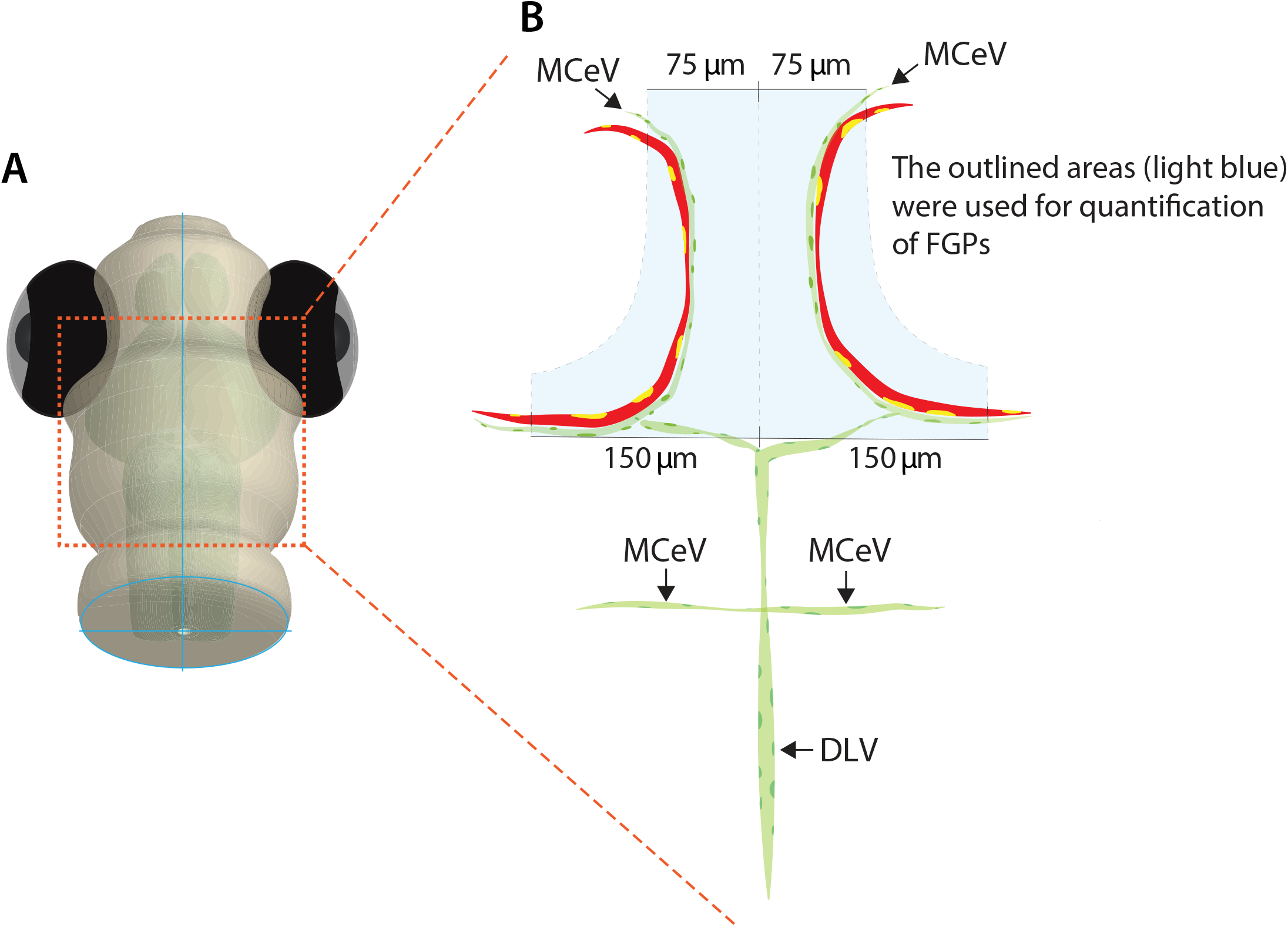
Quantification of FGPs over the optic tectum. (**A**) Schematic representation of the dorsal view of the zebrafish larval head. The boxed area indicates the approximate location where the confocal images of bilateral *Tg(lyve1:*DsRed)^+^ loops over the optic tectum were captured. (**B**) Schematic diagram of the vasculature (green) and bilateral *Tg(lyve1:*DsRed)^+^ loops (red) over the optic tectum. The areas in which FGPs were quantified are highlighted in light blue. The number of *Tg(fli1:*nEGFP*);Tg(lyve1:*DsRed)-double positive FGPs (yellow) within the highlighted area were recorded.

**Figure S2.**
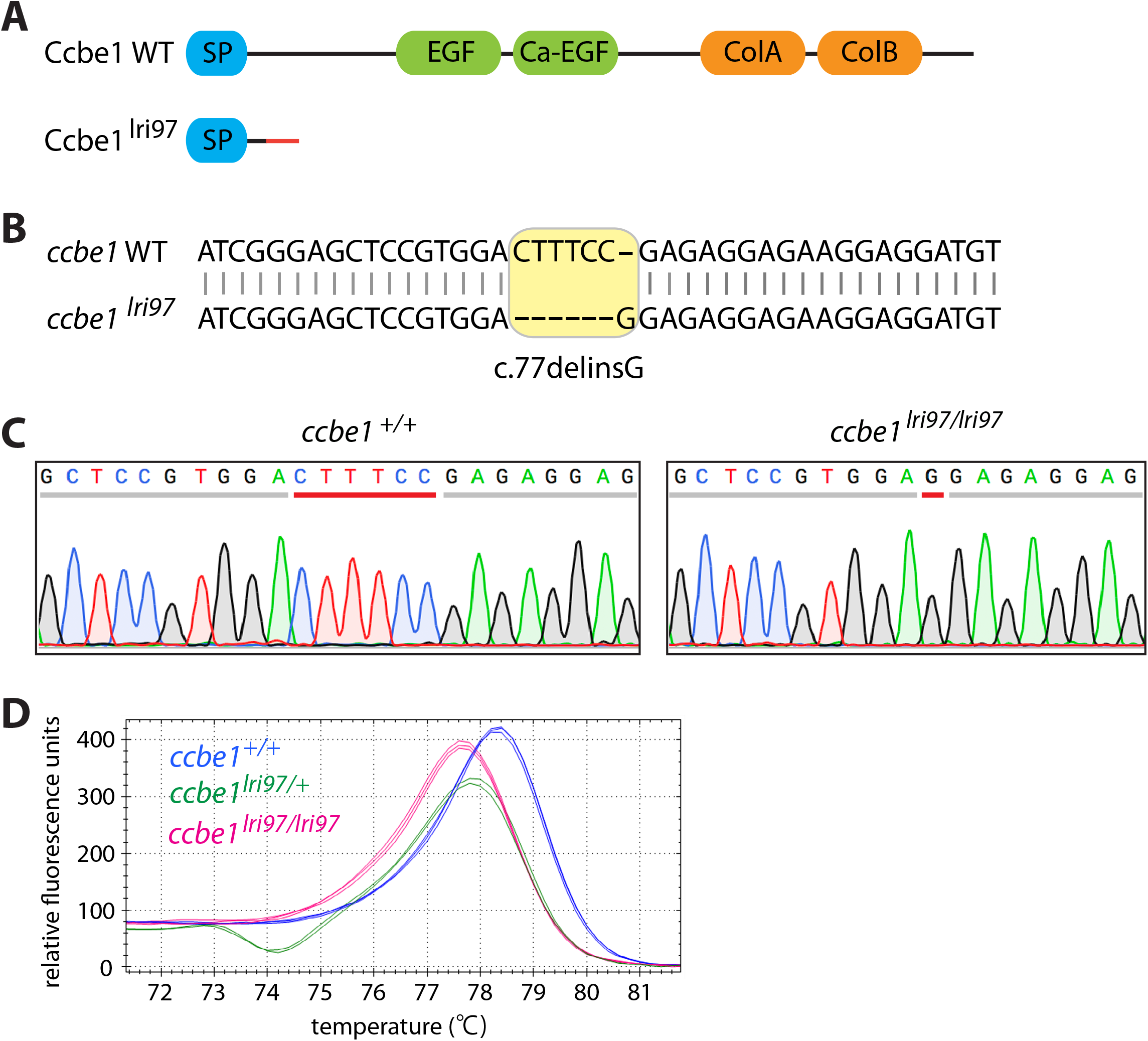
Generation and validation of the *ccbe1^lri97^* mutant allele. (**A**) Predicted protein structure of zebrafish WT Ccbe1 and Ccbe1^lri97^. WT Ccbe1 consists of a signal peptide (SP), an EGF domain (EGF), a calcium-binding EGF domain (Ca-EGF), and two collagen repeat domains (ColA and ColB). Ccbe1^lri97^ is expected to lack all functional domains except a SP. The red solid line of Ccbe1^lri97^ represents missense sequences downstream of the mutation before the stop codon. (**B** and **C**) Sequence alignment of part of exon 1 from the *ccbe1^+/+^* and *ccbe1^lri97/lri97^* allele shows CRISPR-induced indels (**B**). A six-nucleotide deletion (CTTTCC) and a one-nucleotide insertion (G) in the *ccbe1^lri97/lri97^* allele were confirmed by the sequence chromatograms of the PCR products generated using *ccbe1^+/+^* and *ccbe1^lri97/lri97^* fish genomic DNA as templates (**C**). Red lines in the sequence chromatograms indicate the CRISPR-induced nucleotide changes between *ccbe1^+/+^* and *ccbe1^lri97/lri97^* fish (**C**). (**D**) HRMA used to determine the *ccbe1^lri97^* genotypes. Melting curves for 2-3 independent fish of each genotype are shown in different colors as indicated in the panel.

**Figure S3.**
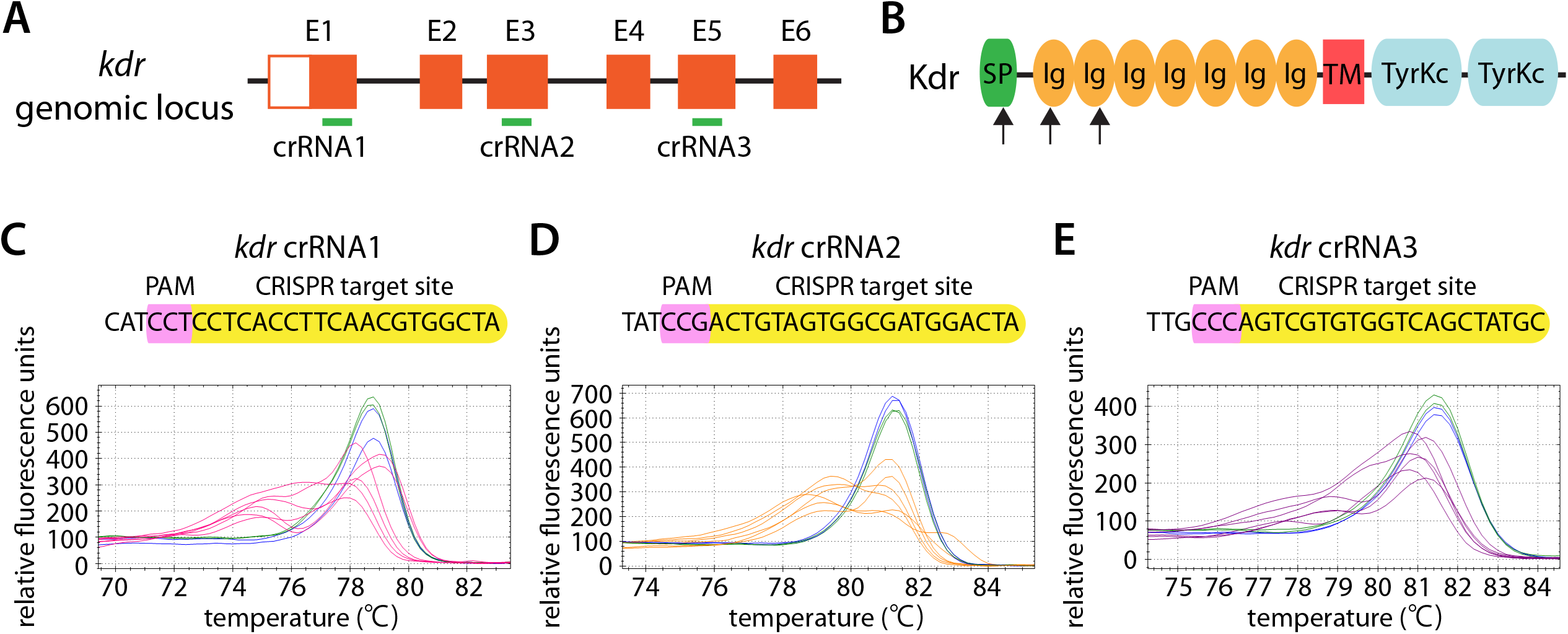
Target design and validation of *kdr* crRNAs. (**A**) Three synthetic crRNAs were designed to target sequences within exon 1 (E1), E3, and E5 on the *kdr* genomic locus. (**B**) Predicted domain structure of zebrafish Kdr. Kdr consists of a signal peptide (SP), seven immunoglobulin-like domains (Ig), a transmembrane domain (TM), and two tyrosine kinase domains (TyrKc). Arrows indicate the approximate positions of the protein sequences corresponding to the target sequences of the three designed crRNAs. (**C**–**E**) HRMA used to validate the efficacy of the three designed crRNAs. The melting curves of 6 independent embryos injected with the dgRNP complex containing *kdr* crRNA1 (**C**, **pink**), crRNA2 (**D**, **orange**), or crRNA3 (**E**, **purple**) are presented. The melting curves of 2 independent, uninjected (**C**–**E**, **blue**) and Cas9-injected (**C**–**E**, **green**) sibling embryos are also presented for comparison for each crRNA. Injection of each dgRNP cocktail disrupted the corresponding target genomic sequences.

**Figure S4.**
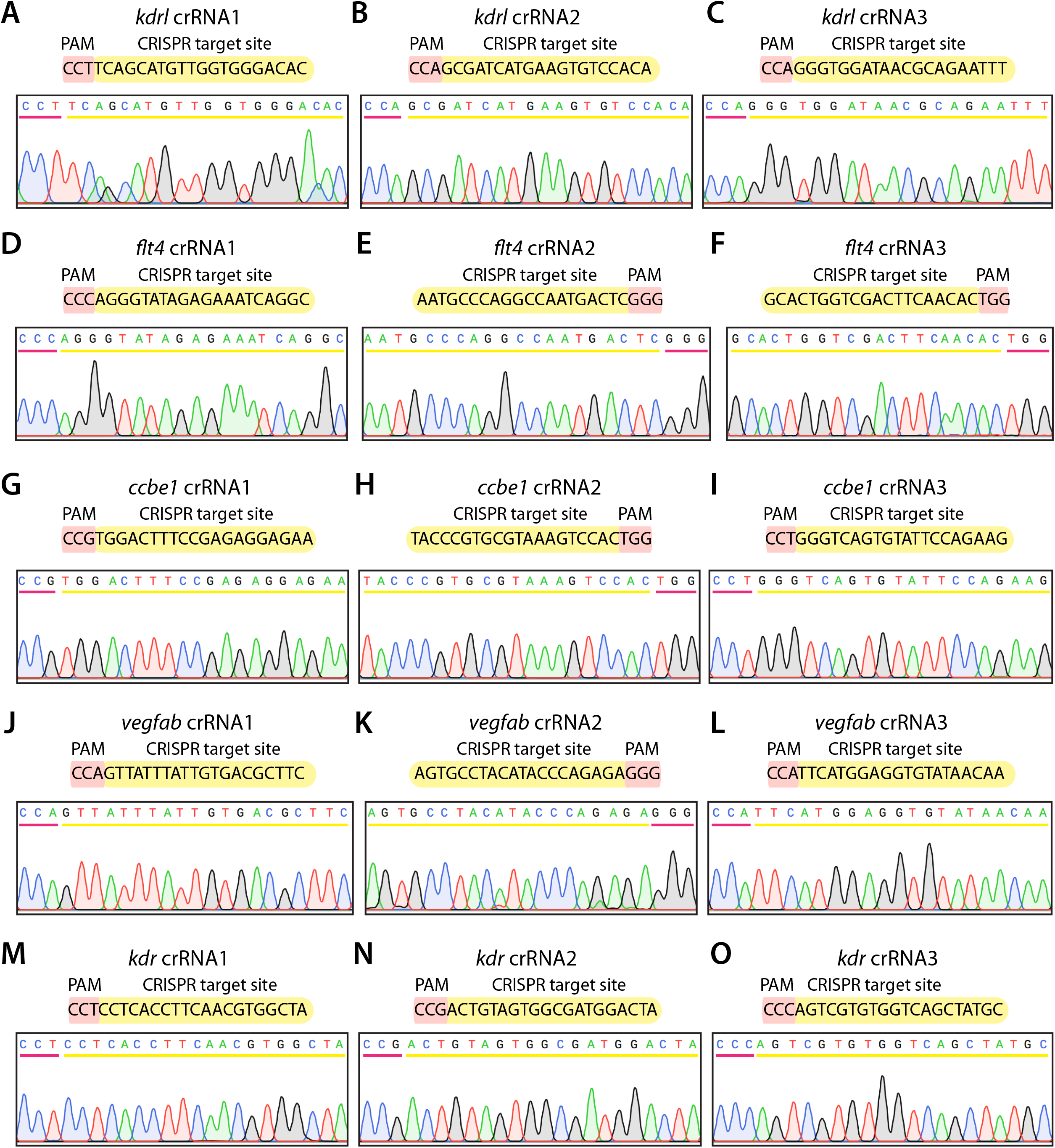
Verification of each designed crRNA’s target sites and PAM sequences within the genome of the fish line used for phenotypic quantifications. The sequence chromatograms of the PCR products that were generated using *Tg(kdrl:EGFP);Tg(kdrl:NLS-mCherry)* (**A**–**C**, **J**–**L**, and **M**–**O**) or *Tg(fli1:*nEGFP*);Tg(lyve1:*DsRed) (**D**–**I**) fish genomic DNA as templates. A sequence chromatogram of the individual crRNA’s target sites (yellow) and PAM sequences (pink) in the genomes of the corresponding fish line is presented. All of the designed crRNAs’ target sites and PAM sequences were perfectly matched to those in the fish lines used for quantifications in this study.

## Notes

### Competing Interest Statement

The authors have declared no competing interest.

